# Metabolomic and transcriptomic analyses of *Fmo5^-/-^* mice reveal roles for flavin-containing monooxygenase 5 (FMO5) in NRF2-mediated oxidative stress, the unfolded protein response, lipid homeostasis, and carbohydrate and one-carbon metabolism

**DOI:** 10.1101/2023.02.09.527806

**Authors:** Ian R. Phillips, Sunil Veeravalli, Sanjay Khadayate, Elizabeth A. Shephard

**Affiliations:** Department of Structural and Molecular Biology, University College London, London, United Kingdom; School of Biological and Chemical Sciences, Queen Mary University of London, London, United Kingdom; MRC London Institute of Medical Sciences (LMS), London, United Kingdom

## Abstract

Flavin-containing monooxygenase 5 (FMO5) is a member of the FMO family of proteins best known for their roles in the detoxification of foreign chemicals and more recently in endogenous metabolism. We have previously shown that *Fmo5^-/-^* mice display an age-related lean phenotype, with much reduced weight gain from 20 weeks of age. The phenotype is characterized by decreased fat deposition, lower plasma concentrations of glucose and cholesterol, higher glucose tolerance and insulin sensitivity, and resistance to diet-induced obesity. In the present study we report the use of metabolomic and transcriptomic analyses of livers of *Fmo5^-/-^* and wild-type mice to identify factors underlying the lean phenotype of *Fmo5^-/-^* mice and gain insights into the function of FMO5. Disruption of the *Fmo5* gene has wide-ranging effects on the abundance of metabolites and expression of genes in the liver. The results reveal that FMO5 is involved in upregulating the NRF2-mediated oxidative stress response, the unfolded protein response and response to hypoxia and cellular stress, indicating a role for the enzyme in adaptation to oxidative and metabolic stress. FMO5 also plays a role in stimulating a wide range of metabolic pathways and processes, particularly ones involved in the regulation of lipid homeostasis, the uptake and metabolism of glucose, the generation of cytosolic NADPH, and in one-carbon metabolism. The results predict that FMO5 acts by stimulating the NRF2, XBP1, PPARA and PPARG regulatory pathways, while inhibiting STAT1 and IRF7 pathways.

## Introduction

Mammalian flavin-containing monooxygenases (FMOs) (EC 1.14.13.8) are a small family of proteins (FMOs 1-5) whose members play roles in both xenobiotic [1] and endogenous metabolism [2]. FMOs catalyze the NADPH-dependent oxygenation of a wide range of chemicals. Drug substrates of FMO1 and FMO3 can overlap and are mono-oxygenated usually at a nitrogen or a sulfur atom [1]. One of the substrates of FMO3 is the dietary-derived odorous chemical trimethylamine [3], which is converted in the liver to its non-odorous *N*-oxide. Mutations in the human *FMO3* gene cause the inherited disorder trimethylaminuria [4]. FMO5 is unusual in that its substrates do not overlap with those of other FMOs and it catalyzes Baeyer-Villiger reactions [5,6].

The use of *Fmo* knockout mouse lines has been effective in establishing functions for FMOs. FMO1 has been identified as a key participant in the metabolism of drugs subject to retro-reduction [7], as a regulator of energy balance [8] and as an enzyme that converts hypotaurine to taurine [9]. Male mice are natural liver-specific knockouts for FMO3 [10], with *Fmo3* expression being switched off at about 6 weeks of age [11], and consequently excrete large amounts of trimethylamine in their urine.

We have previously shown that *Fmo5^-/-^* mice display an age-related lean phenotype, with much reduced weight gain from about 20 weeks of age [12]. The phenotype is characterized by decreased fat deposition, lower plasma concentrations of glucose and cholesterol, higher glucose tolerance and insulin sensitivity, and resistance to diet-induced obesity [12,13]. FMO5 is expressed in the liver and throughout the gut [13] and acts as a potential sensor of gut bacteria [13]. In many respects the phenotype of *Fmo5^-/-^*mice is similar to that of germ-free mice [13,14], supporting a role for FMO5 in host-microbiome interactions. Treatment of *Fmo5^-/-^*mice with antibiotics does not change their phenotype with respect to plasma glucose, glucose tolerance or insulin sensitivity. In contrast, wild-type (WT) mice treated with antibiotics showed improved glucose tolerance and insulin sensitivity [13]. Analysis of gut microbiome composition [13] and the presence of the microbial product 2,3 butanediol in the urine of *Fmo5^-/-^*, but not of WT mice [15] confirms that FMO5 plays a role in modulating microbial gut populations. Expression of recombinant mouse FMOs 1, 2, 3, 4 and 5 in cell lines derived from kidney or liver showed that each of the proteins affected metabolic processes and protected against chemically induced stress [16].

FMO5 is highly expressed in liver in both mouse and human [11,17,18]. In the present study we report the use of metabolomic and transcriptomic analyses of livers of *Fmo5^-/-^*and WT mice to gain insights into factors underlying the lean phenotype of *Fmo5^-/-^*mice. The results indicate that FMO5 plays a role in the NRF2-mediated oxidative stress response, the unfolded protein response and in several metabolic pathways and processes, particularly ones involved in the regulation of lipid homeostasis, the uptake and metabolism of glucose, the generation of cytosolic NADPH, and in one-carbon metabolism.

## Results

Transcriptomic analysis of 15,570 genes, with cut-offs of p_adj_(q value) < 0.05 and base mean > 500, found that 744 genes were differentially expressed in the liver of *Fmo5^-/-^* mice compared with that of WT mice (Fig 1A). The expression of 216 was increased (ranging from 1.2- to 80-fold) and that of 528 was decreased (by 16 to 98.5%) in *Fmo5^-/-^*mice.

**Fig 1.**
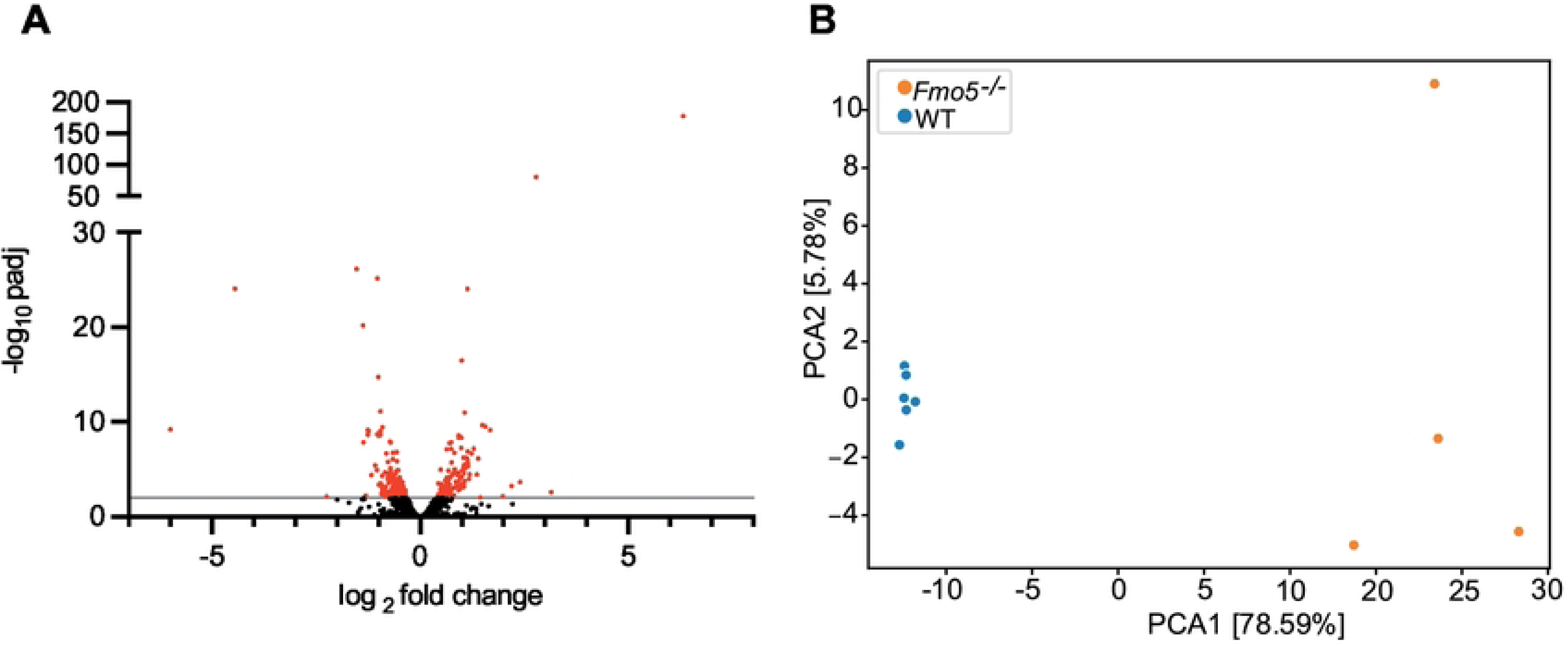
Transcriptomic and metabolomic analyses of *Fmo5^-/-^* and WT mice. (A) Volcano plot representation of differentially expressed genes (log_2_-fold change) of *Fmo5^-/-^* vs WT mice. Red dots are genes whose expression is significantly increased or decreased in *Fmo5^-/-^* mice with a -log_10_ padj above 2. (B) PCA analysis of the liver metabolome of *Fmo5^-/-^* vs WT mice.

Metabolomic analysis of mouse liver identified 686 compounds (S1 Table). The abundance of 318 differed significantly (*p* < 0.05) between *Fmo5^-/-^*and WT mice. Of these, 202 were more abundant (from 1.03- to 22-fold) and 116 less abundant (from 4.7 to 95%) in *Fmo5^-/-^*mice. Principal component analysis (PCA) of hepatic metabolites confirmed that *Fmo5^-/-^*and WT mice differed significantly, with PC1 explaining 78.59% of the total difference (Fig 1B).

Analysis of the transcriptomic dataset by IPA software revealed that in the liver of *Fmo5^-/-^* mice the major downregulated canonical pathways were NRF2-mediated oxidative stress response (*p* = 4.30e-12, z score −2.138) and the unfolded protein response (*p* = 3.46e-10, z score −3.051), and the major downregulated metabolic functions involved transport and metabolism of lipids (Fig 2).

**Fig 2.**
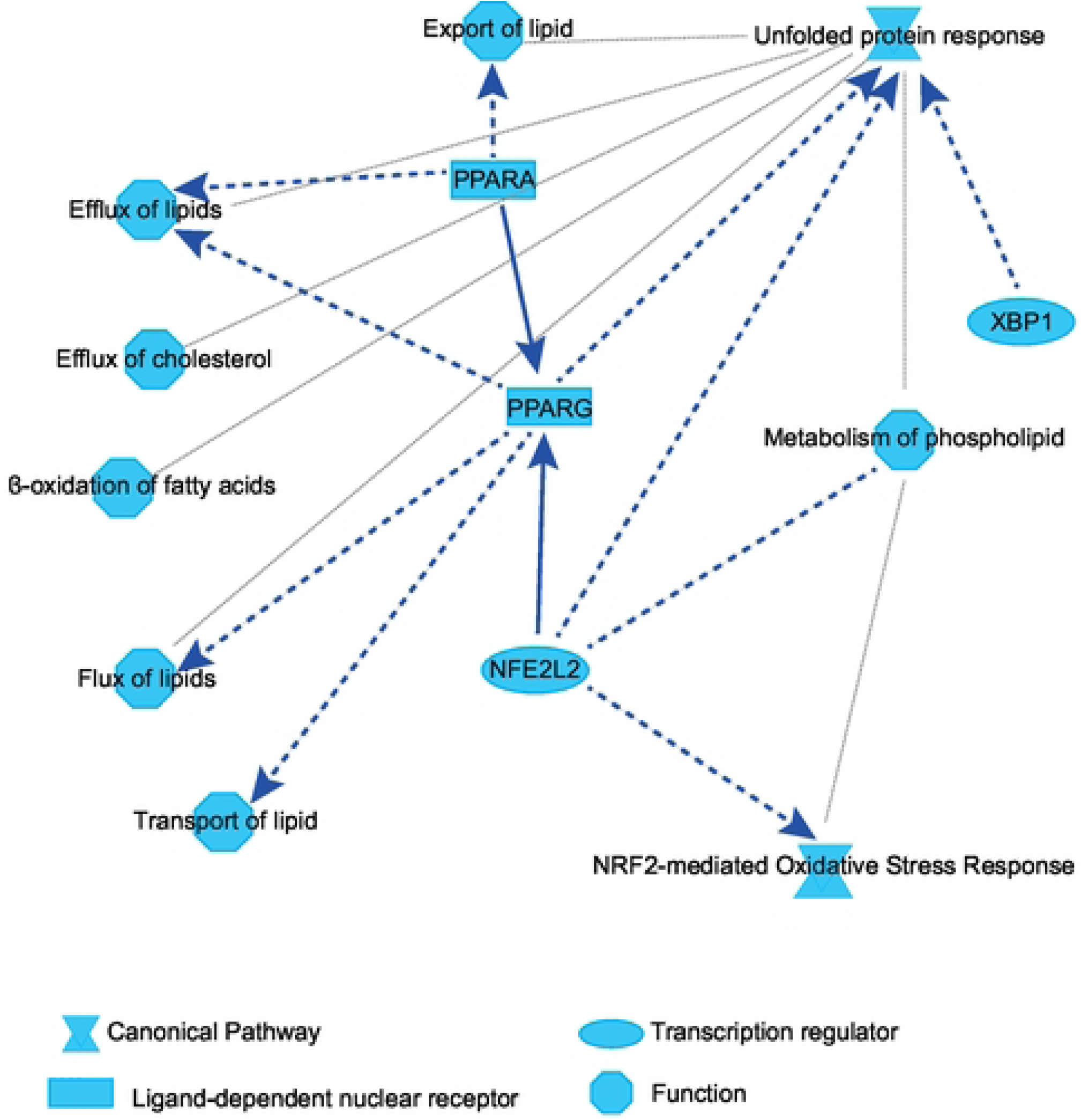
Graphical summary of relationships predicted by IPA to be down-regulated in *Fmo5*^-/-^ mice. Solid line: direct interaction; dashed line: indirect interaction; grey dotted line: inferred correlation from machine-based learning.

### NRF2-mediated oxidative stress response

The expression in liver of *Nfe2l2*, which encodes NRF2, was lower in *Fmo5^-/-^* than in WT mice (Table 1). NRF2-induced genes that are down-regulated in liver of *Fmo5^-/-^* mice (Table 1) include those encoding the detoxification enzymes FMO1, microsomal epoxide hydrolase 1 (EPHX1), microsomal glutathione S-transferase 1 (MGST1), aldehyde oxidase 1 (AOX1) and several UDP glucuronosyltransferases and carboxylesterases.

**Table 1.** NRF2-mediated oxidative stress genes differentially expressed in liver of Fmo5^-/-^ mice.

| Role | Gene symbol | Protein | Percentage change | padj |
| --- | --- | --- | --- | --- |
| **Detoxification** | *Aox1* | Aldehyde oxidase 1 | -30 | 0.036 |
|  | *Ces1d* | Carboxylesterase 1D | -29 | 0.050 |
|  | *Ces1e* | Carboxylesterase 1E | -40 | 1.08e-4 |
|  | *Ces1f* | Carboxylesterase 1F | -24 | 0.035 |
|  | *Ephx1* | Epoxide hydrolase 1 | -33 | 1.62e-5 |
|  | *Fmo1* | Flavin-containing monooxygenase 1 | -24 | 0.006 |
|  | *Mgst1* | microsomal glutathione S-transferase 1 | -27 | 0.001 |
|  | *Ugt1a6b* | UDP glucuronosyltransferase 1A6B | -37 | 0.001 |
|  | *Ugt2b5* | UDP glucuronosyltransferase 2B5 | -36 | 0.001 |
|  | *Ugt2b34* | UDP glucuronosyltransferase 2B34 | -32 | 0.001 |
|  | *Ugt2b36* | UDP glucuronosyltransferase 2B36 | -28 | 0.004 |
|  | *Ugt3a1* | UDP glucuronosyltransferase 3A1 | -39 | 0.005 |
| **Glutathione synthesis** | *Gclc* | Glutamate-cysteine ligase, catalytic subunit | -36 | 0.022 |
|  | *Gclm* | Glutamate-cysteine ligase, modifier subunit | -22 | 0.028 |
| **Glutathione utilization** | *Prdx1* | Peroxiredoxin 1 | -21 | 0.024 |
|  | *Prdx3* | Peroxiredoxin 3 | -16 | 0.040 |
| **NADPH regeneration** | *Idh1* | Isocitrate dehydrogenase (NADP<sup>+</sup>) 1 (cytosolic) | -29 | 8.20e-4 |
|  | *Me1* | Malic enzyme 1 (cytosolic) | -60 | 0.006 |
| **Regulators** | *Maf* | MAF BZIP transcription factor | +42 | 8.02e-4 |
|  | *Nfe2l2* | Nuclear factor, erythroid 2 like 2 (NRF2) | -19 | 0.033 |

Many products of phase-1 detoxification reactions are conjugated with glutathione. Expression of *Gclc* and *Gclm*, NRF2-activated genes that encode the two components of glutamate-cysteine ligase, which catalyzes the rate-limiting step in the synthesis of glutathione, was lower in *Fmo5^-/-^*mice. Glutathione is also required for reduction of H_2_O_2_, in reactions catalyzed by glutathione peroxidase or peroxiredoxins (PRDXs), and expression of *Prdx1* and *3* was lower in *Fmo5^-/-^*mice. Many detoxification reactions require NADPH, and expression of the NRF2-activated genes *Idh1* and *Me1*, encoding isocitrate dehydrogenase 1 and cytosolic malic enzyme, which catalyze the regeneration of NADPH, was lower in *Fmo5^-/-^* mice. The abundance of ME1 protein also is lower in liver of *Fmo5^-/-^* mice [12].

NRF2 activates transcription by binding to antioxidant response elements (AREs). The expression of *Maf*, which encodes a protein that represses ARE-mediated transcription [19], was higher in *Fmo5^-/-^* mice (Table 1), and would thus contribute to the suppression of NRF2-mediated oxidative stress response in these animals.

### The unfolded protein response (UPR)

The expression of several genes encoding proteins involved in protein folding, modification and quality control in the cytosol, the endoplasmic reticulum (ER) or mitochondria was lower in *Fmo5^-/-^* mice (Table 2): *Canx*, *Cct5*, *Cct6a*, *Cct8*, encoding chaperones, genes encoding chaperones of the heat-shock protein (HSP) 70 family, including *Hspa5* (BiP), one of the most significantly differentially expressed genes (padj = 8.01e-12), and of the HSP90, HSP110, HSP60 and HSP40 families; genes *Stip1*, *St13* and *Fkbp4,* encoding co-chaperones; *Pdia3*, *Pdia4*, *Pdia6* and *Erp44*, encoding protein disulfide-isomerases that promote protein folding through formation of disulfide bonds and act as chaperones to prevent aggregation of misfolded proteins; *Serp1*, encoding a protein that protects unfolded proteins against degradation during ER stress; *Uggt1*, which encodes a UDP-glucose:glucosoyltransferase that catalyzes the selective reglucosylation of unfolded glycoproteins; and *Mlec*, encoding malectin, involved in regulating glycosylation in ER. Protein disulfide-isomerases are reoxidized by ER oxidoreductin, in a process that generates reactive oxygen species (ROS). Expression of *Ero1b*, encoding the beta isoform of ER oxidoreductin, which is induced by the UPR [20], was lower in *Fmo5^-/-^* mice (Table 2). The results suggest that the UPR is attenuated in the liver of *Fmo5^-/-^* mice.

**Table 2.**
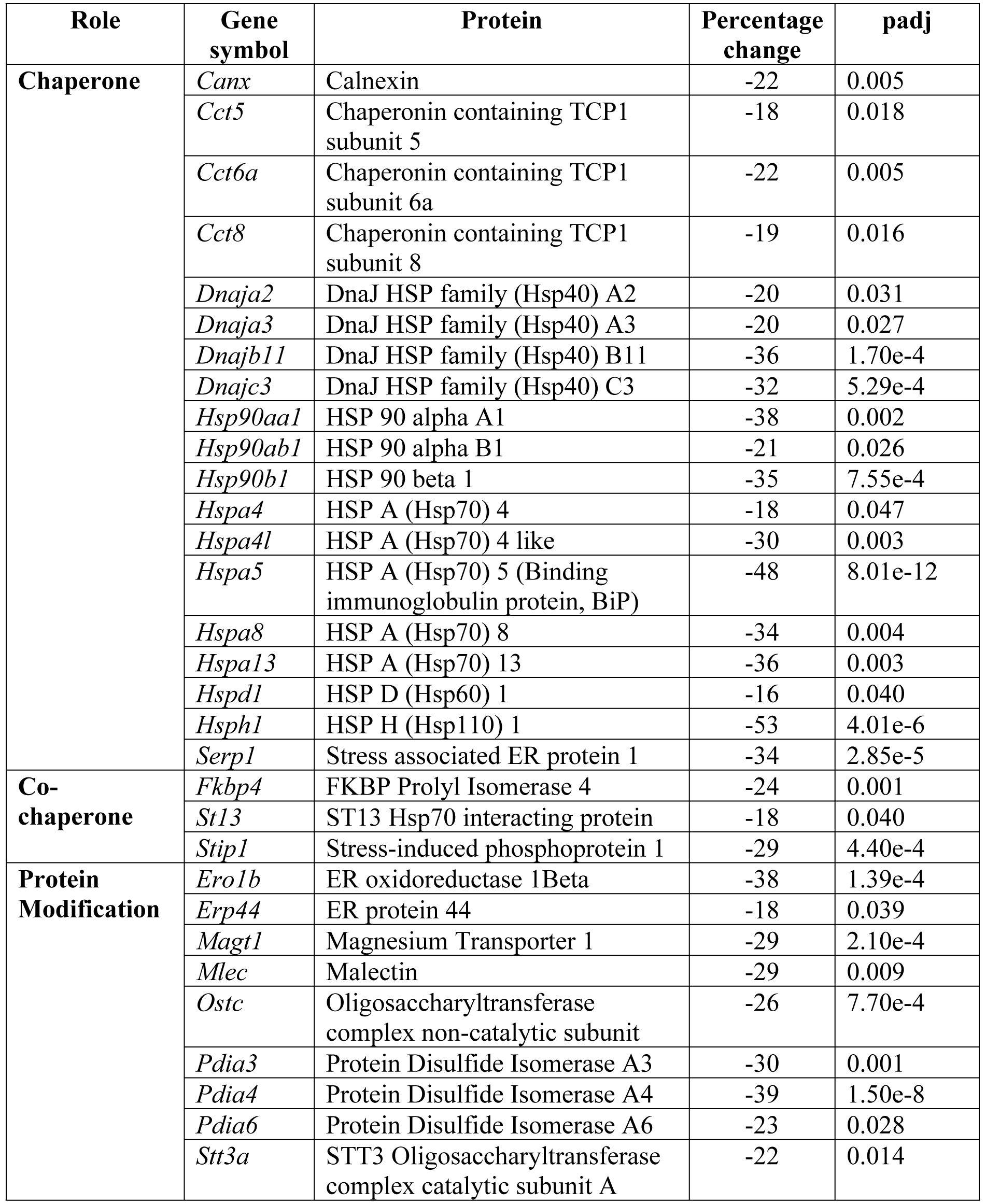
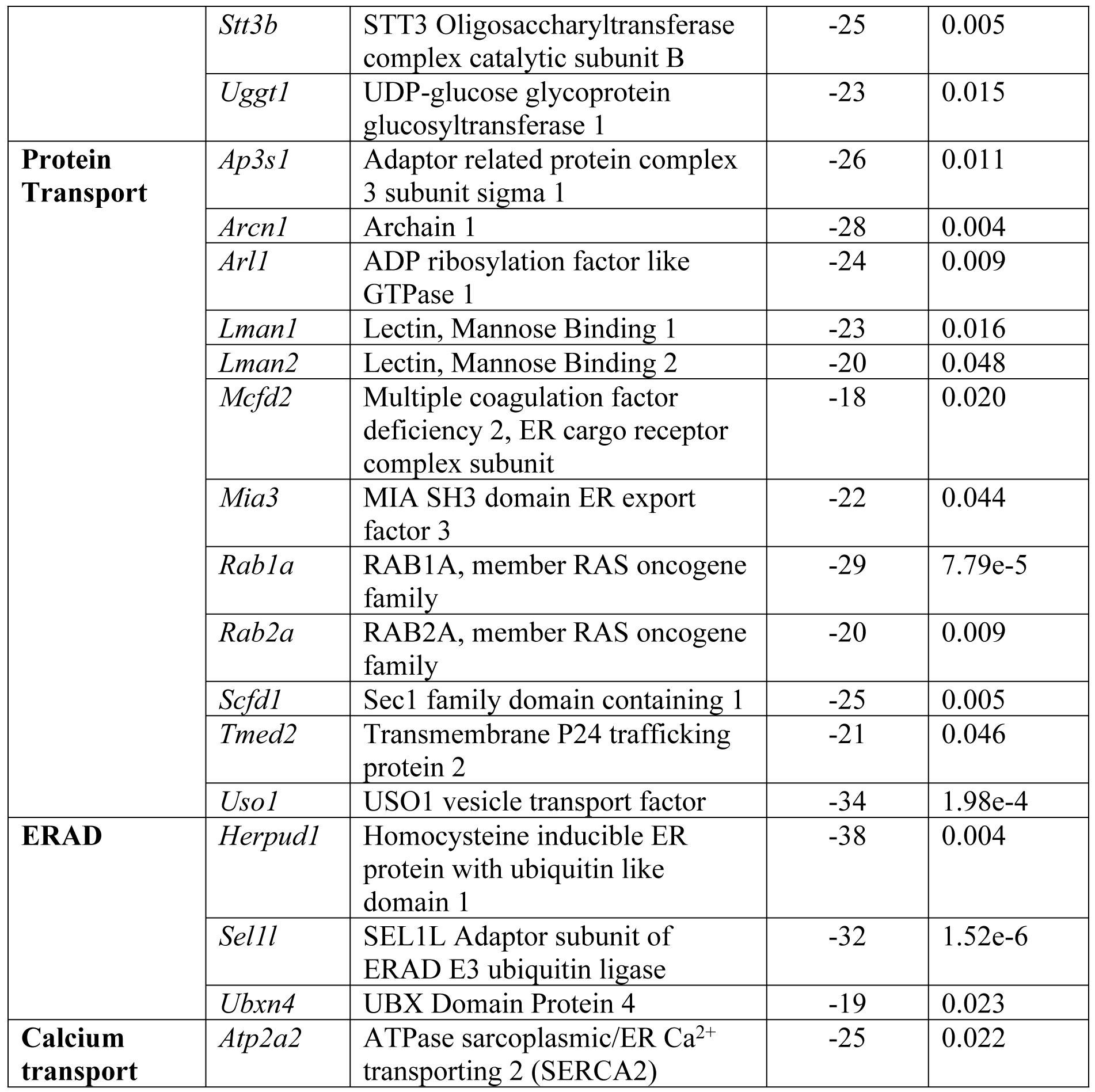
Expression of genes encoding proteins involved in protein folding, quality control, trafficking and ER-associated degradation differentially expressed in liver of *Fmo5^-/-^* mice.

HSPA5 (BiP) and HSP90B1 also act to recruit misfolded proteins to the ER-associated degradation (ERAD) pathway [21,22]. The lower expression of *Hspa5* and *Hsp90b1*, together with that of *Herpud1*, encoding a UPR-induced adaptor specific for degradation of non-glycosylated proteins, *Sel1l*, encoding an adaptor required for the stability of HRD1[23], a key ubiquitin E3 ligase component of the ERAD pathway, and *Ubxn4* (Table 2), encoding a protein which promotes ERAD, suggests that the ERAD is attenuated in *Fmo5^-/-^*mice.

The oligosaccharide transferase (OST) complex, located in the membranes of the ER, catalyzes the first step in protein glycosylation, the addition of *N*-acetylglucosamine to a nascent protein. The expression of *Stt3a* and *Stt3b*, encoding the two catalytic subunits of OST, and that of *Magt1* and *Ostc*, encoding two other components of the complex, was lower in *Fmo5^-/-^*mice (Table 2). The expression of several genes encoding proteins involved in vesicular protein trafficking also was lower in *Fmo5^-/-^*mice: *Lman1*, *Lman2*, *Arcn1*, *Mcfd2*, *Mia3*, *Rab1a*, *Rab2a* and *Scfd1*, involved in transport from the ER to the Golgi; *Uso1*, in transport within the Golgi; *Ap3s1*, in transport from the Golgi; and *Arl1* and *Tmed2*, in general vesicular transport (Table 2), suggesting that vesicular transport is perturbed in *Fmo5^-/-^* mice.

The UPR also increases ER calcium stores, triggering calcium-dependent inflammatory responses. Expression of *Atp2a2*, encoding SERCA2, which transports Ca^2+^ into the ER, was lower in *Fmo5^-/-^* mice (Table 2), suggesting that *Fmo5^-/-^* mice have lower levels of inflammation. Consistent with this, *Fmo5^-/-^* mice have lower amounts of inflammatory markers [13].

### Lipid metabolism

There were marked differences between *Fmo5^-/-^* and WT mice in the hepatic concentrations of several lipids (S1 Table, Fig 3A). In addition, several genes encoding proteins involved in lipid metabolism were differentially expressed in the livers of *Fmo5^-/-^* and WT mice (Table 3) and IPA identified several functions of lipid metabolism that were perturbed in *Fmo5^-/-^* mice (Fig 2).

**Table 3.**
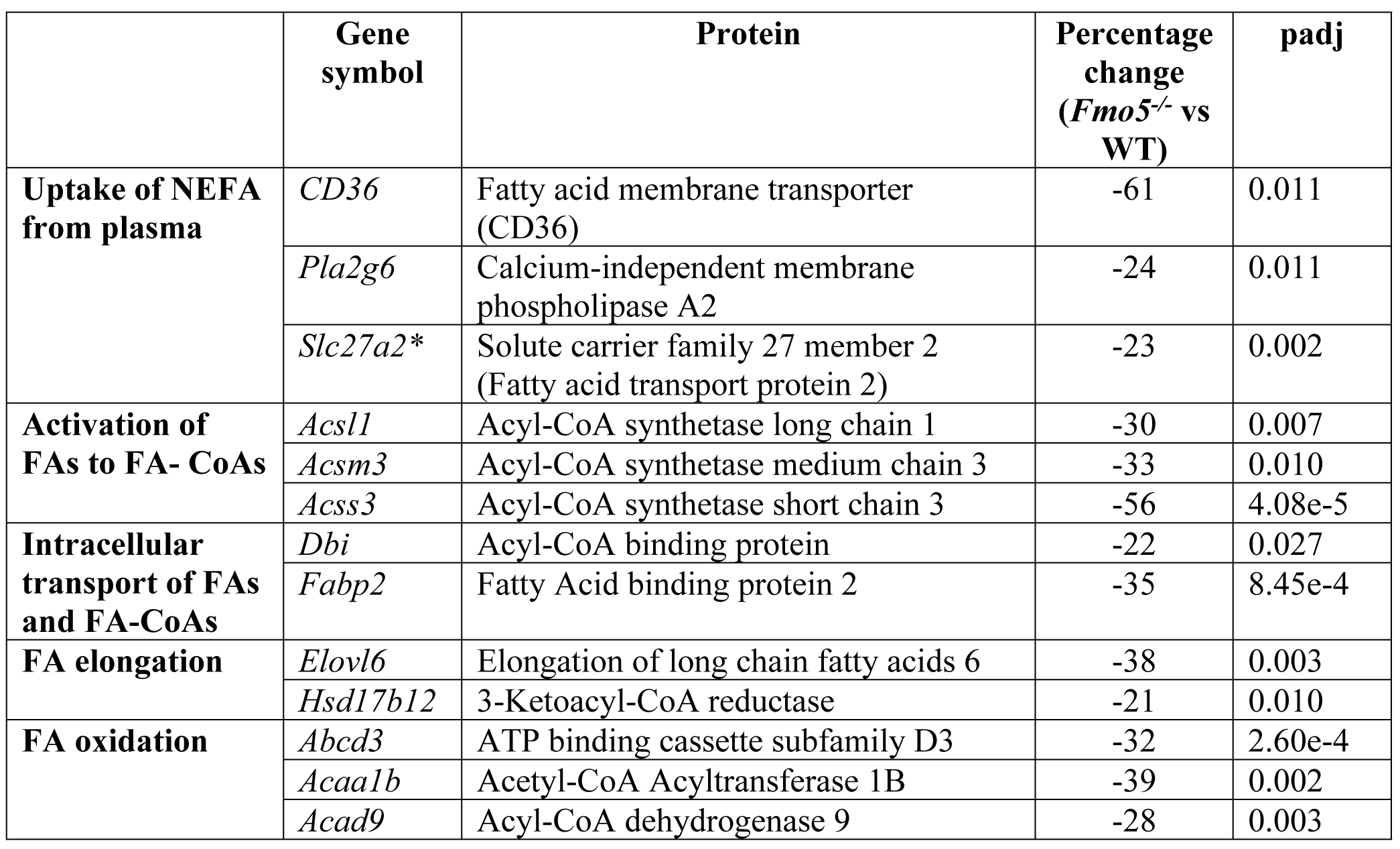
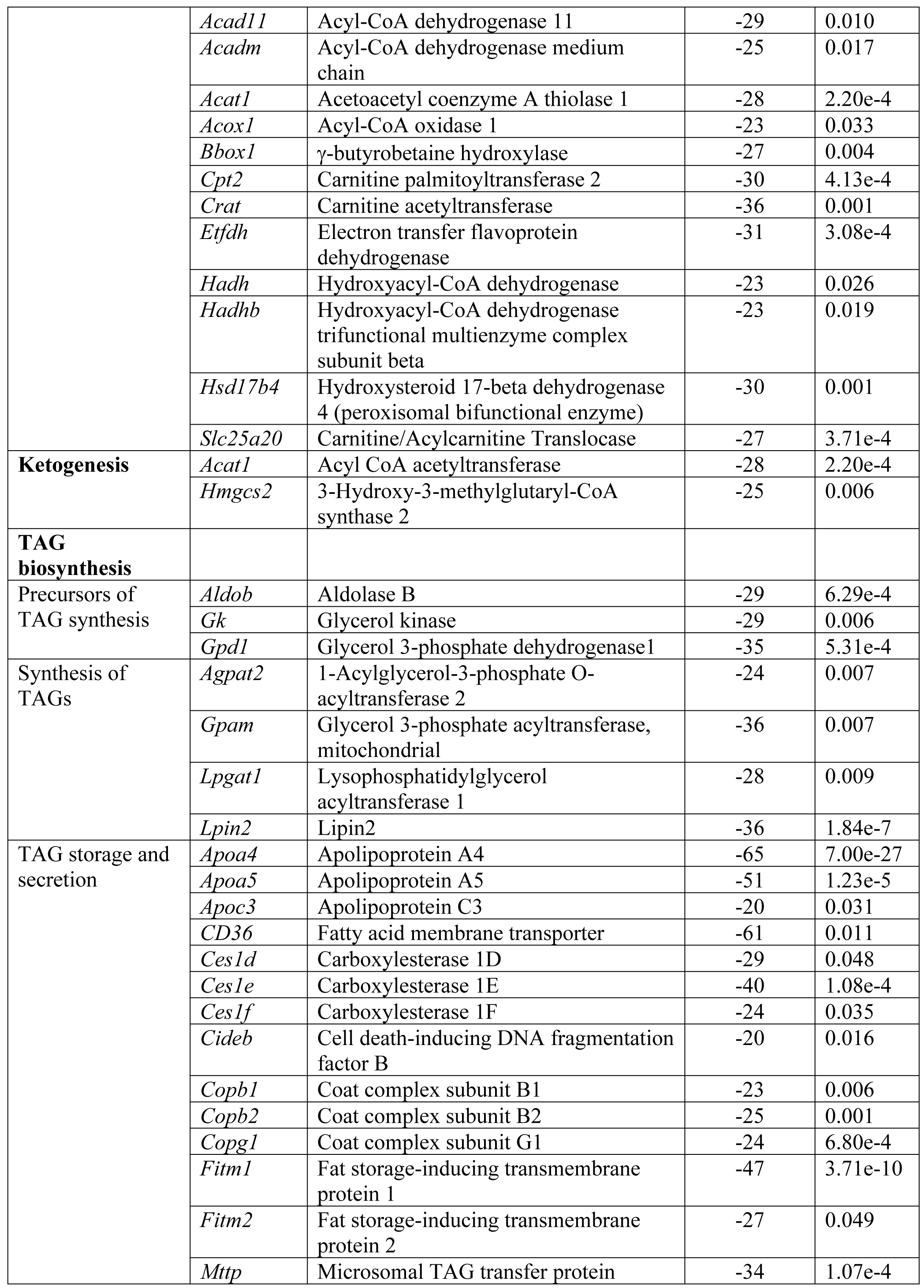
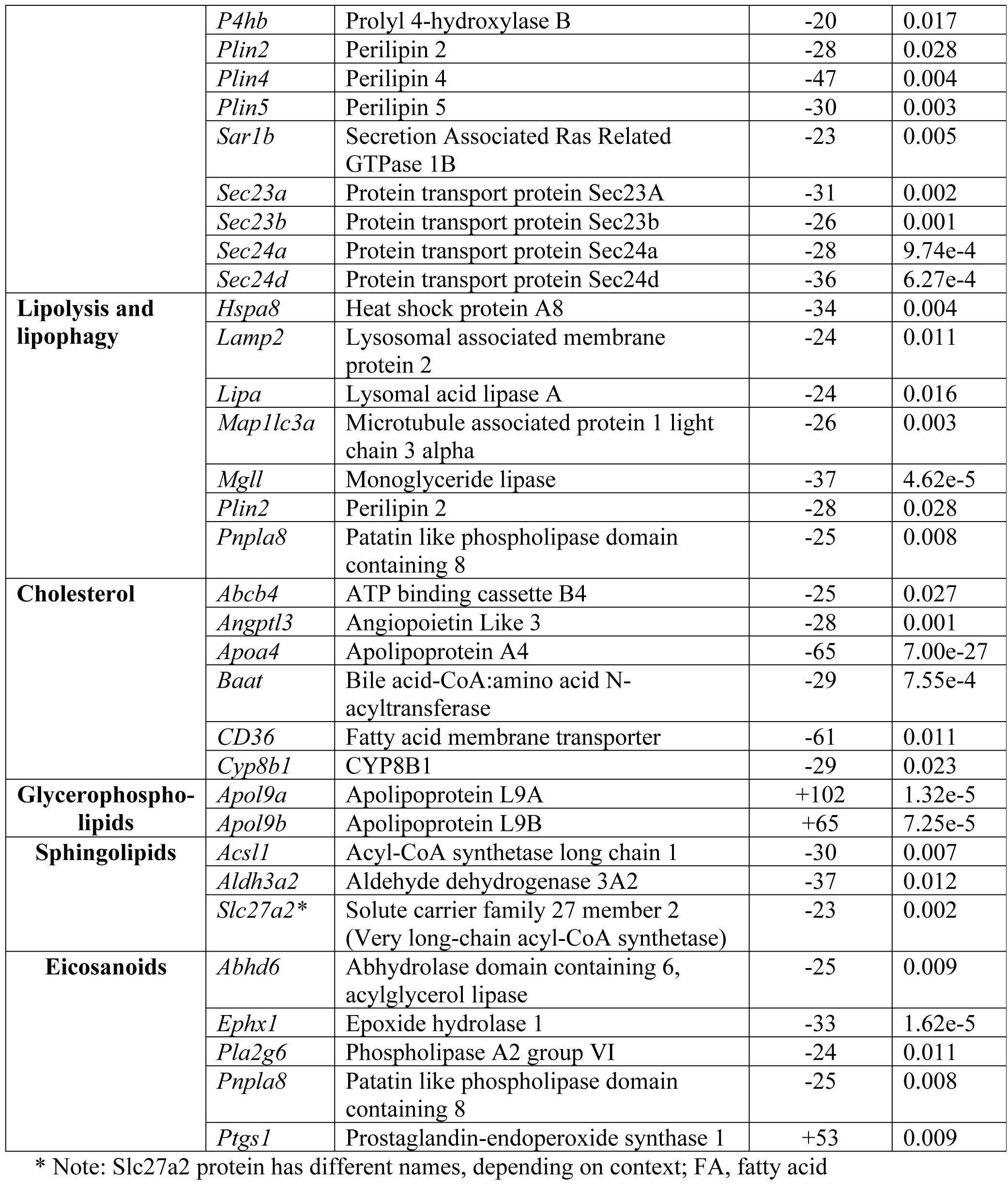
Genes involved in lipid metabolism that are differentially expressed in the liver of *Fmo5^-/-^* mice.

**Fig 3.**
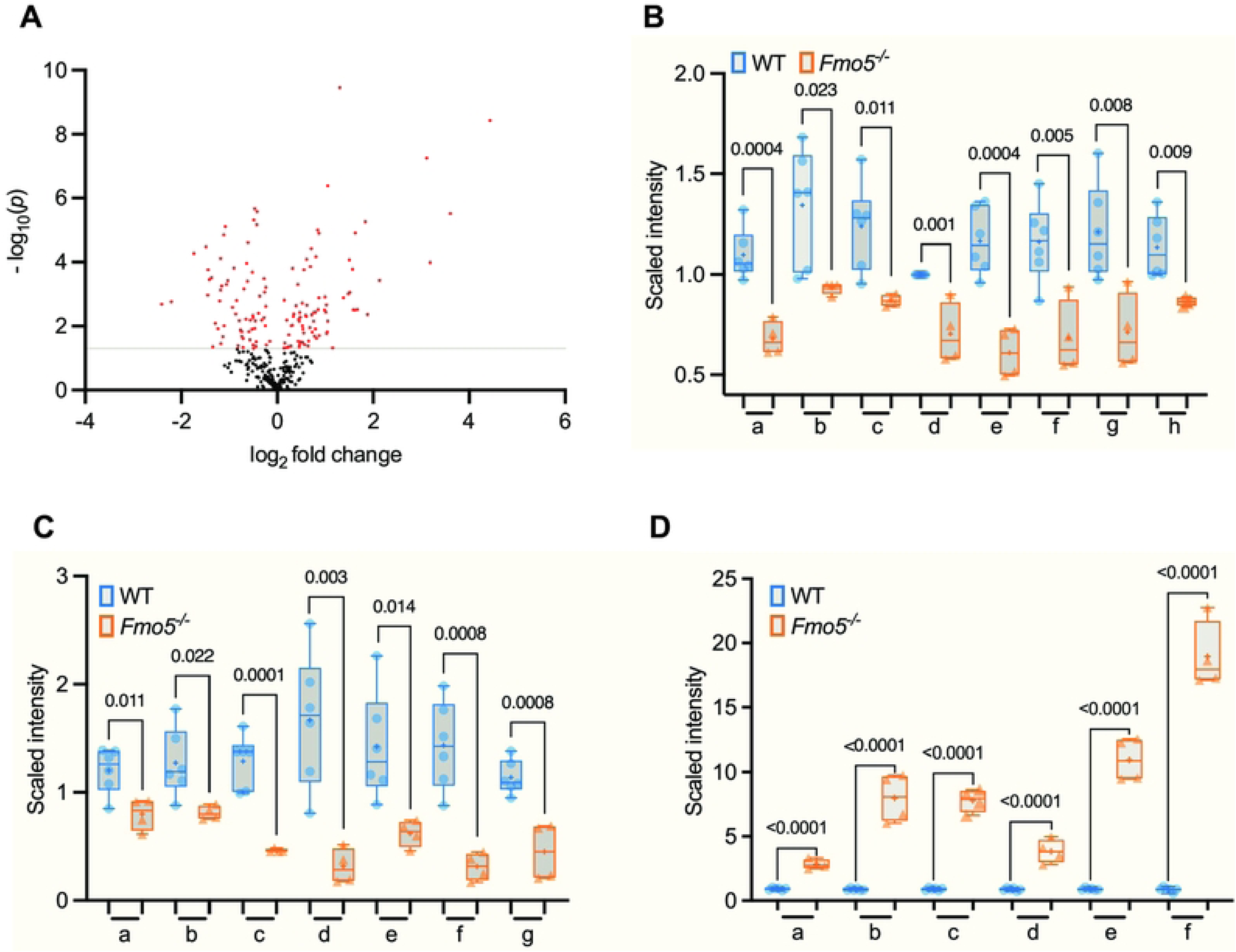
Differences in abundance of lipid molecules between *Fmo5^-/-^* vs WT mice. (A) Volcano plot representation of lipid molecules (log_2_-fold change) of *Fmo5^-/-^* vs WT mice. Red dots are lipids that are significantly increased or decreased in *Fmo5^-/-^* mice with a -log_10_ (A) (*p*) above 1.301 (grey line). (B) Long-chain saturated and monounsaturated lipids: a. myristate (14:0); b. nonadecanoate (19:0); c. arachidate; d. palmitoleate (16:1n7); e. 10-heptadecenoate (17:1n7); f. 10-nonadecenoate (19:1n9); g. eicosenoate (20:1); h. margarate. (C) Carnitine and long-chain carnitines: a. carnitine; b. stearoylcarnitine (C18); c. arachidoylcarnitine (C20)*; d. behenoylcarnitine (C22)*; e. lignoceroylcarnitine (C24)*; f. erucoylcarnitine (C22:1)*; g. nervonoylcarnitine (C24:1)*. (D) Complex lipids and sphingolipids: a. sphinganine; b. N-palmitoyl-sphinganine (d18:0/16:0); c. palmitoyl dihydrosphingomyelin (d18:0/16:0)*; d. behenoyl dihydrosphingomyelin (d18:0/22:0)*; e. sphingomyelin (d18:0/18:0, d19:0/17:0)*; f. sphingomyelin (d18:0/20:0, d16:0/22:0)*. + indicates mean value, numbers above box plots are *p* values.

### Uptake of non-esterified fatty acids (NEFA) from plasma

The hepatic expression of *Cd36*, a gene that is activated by NRF2, and *pla2g6*, both of which encode proteins involved in fatty acid uptake [24], was lower in *Fmo5^-/-^* mice than in WT mice, as was that of the gene encoding SLC27A2 (Table 3), which couples the uptake of fatty acids with their intracellular esterification into acyl-CoAs. The decreased expression of fatty acid transporters suggests that, although the plasma concentration of NEFA in *Fmo5^-/-^*and WT mice was similar (S2 Table), as was found previously [12], in *Fmo5^-/-^*mice uptake of NEFA into liver is compromised. Another source of intracellular fatty acids is de novo biosynthesis. The expression of none of the genes encoding enzymes of the major fatty acid biosynthetic pathway, located in the cytosol, was affected in *Fmo5^-/-^*mice. Expression of genes encoding low-density lipoprotein receptor (LDLR) and LDLR-related protein-1, which are involved in the uptake of chylomicron remnants, was similar in *Fmo5^-/-^*and WT mice, indicating that uptake of triacylglycerol (TAG) from plasma is unaffected in *Fmo5^-/-^*mice.

### Activation of fatty acids to fatty acyl-CoAs

For incorporation into complex lipids or transport into mitochondria for β-oxidation, fatty acids are esterified to CoA to form fatty acyl-CoAs, in reactions catalyzed by acyl-CoA synthetases (Acs). The expression of *Acsl1*, *Acsm3* and *Acss3*, encoding proteins that have substrate preferences for long-, medium- and short-chain fatty acids, respectively, was lower in *Fmo5^-/-^* mice, whereas that of the gene encoding acyl-CoA thioesterases, which catalyze the reverse reaction, fatty acyl-CoA to fatty acid, was similar in *Fmo5^-/-^* and WT mice, suggesting that in *Fmo5^-/-^* mice the balance between fatty acids and acyl-CoAs is shifted towards fatty acids, with less acyl-CoAs being available for TAG synthesis or β-oxidation.

### Intracellular transport of fatty acids and fatty acyl-CoAs

In the cytosol, fatty acids are bound to sterol carrier protein-2 (SCP2), fatty acid binding protein 1 (FABP1) and FABP2, which preferentially binds saturated long-chain fatty acids, whereas fatty acyl-CoAs bind to SCP2 and acyl-CoA binding protein (encoded by the *Dbi* gene). Expression of *Fabp1* and *Scp2* was similar in *Fmo5^-/-^* and WT mice, but that of *Fabp2* and *Dbi* was lower in *Fmo5^-/-^* mice (Table 3). The lower expression of *Fabp2* is consistent with the lower hepatic concentrations of long-chain saturated fatty acids in *Fmo5^-/-^* mice (S1 Table and see below), whereas the lower expression of *Dbi*, in conjunction with similar expression of *Fabp1* and *Scp2*, supports an increased ratio of fatty acids to fatty acyl-CoAs in these mice.

### Fatty acid elongation

Fatty acids are elongated (chain length extended beyond C16) and double-bonds introduced by enzymes in the ER. The elongation cycle consists of four steps. The first, rate-limiting step, a condensation reaction, is catalyzed by ELOVL enzymes, each of which has a substrate preference for different length chains. The expression of *Elovl6*, encoding an enzyme that has a preference for C16 chains and, hence, is involved in addition of the first two carbons in the elongation process, was lower in *Fmo5^-/-^*mice than in WT mice (Table 3). The expression of *Hsd17b12*, encoding an enzyme that catalyzes the second, reduction, step of the cycle, also was lower in *Fmo5^-/-^*mice. The results suggest that elongation of fatty acids beyond a C16 length would be compromised in *Fmo5^-/-^*mice.

### Intracellular hepatic fatty acid concentrations

The hepatic concentrations of eight of the 10 long-chain saturated and monounsaturated fatty acids measured were lower in *Fmo5^-/-^* than in WT mice (S1 Table and Fig 3B). In contrast, the concentrations of only two of 16 long-chain polyunsaturated fatty acids were lower in *Fmo5^-/-^* mice. Most of the long-chain polyunsaturated fatty acids were omega-3 or omega-6 fatty acids, derived from the essential fatty acids linolenic acid (C18:3, n-3) and linoleic acid (C18:2, n-6) by elongation reactions not involving ELOVL6. The concentrations of long-chain saturated and monounsaturated acyl-carnitines, which are in equilibrium with their CoA derivatives, were also lower in the liver of *Fmo5^-/-^* mice (S1 Table), suggesting a decline in β-oxidation. In contrast, concentrations of acyl-cholines were higher in *Fmo5^-/-^* mice. The lower hepatic concentrations of long-chain saturated and monounsaturated fatty acids in *Fmo5^-/-^* mice is consistent with the lower expression in these mice of genes encoding fatty acid elongation enzymes.

### Fatty acid oxidation

Long-chain fatty acyl-CoAs are transferred into the mitochondria for β-oxidation via the carnitine shuttle. In the cytosol, the acyl group of a fatty acyl-CoA is transferred to carnitine in a reaction catalyzed by carnitine acyl transferase 1 (CPT1). Acyl-carnitine is then transported across the inner mitochondrial membrane by carnitine/acylcarnitine translocase (SLC25A20), an antiporter which exchanges fatty acyl carnitine for carnitine. In the mitochondrial matrix the acyl group is then transferred from carnitine to CoA, by CPT2. Hepatic concentrations of carnitine and several long-chain acyl-carnitines were lower in *Fmo5^-/-^* mice (S1 Table and Fig 3C). Carnitine is derived from the diet, but can also be synthesized in vivo, and expression of *Bbox1*, encoding γ-butyrobetaine hydroxylase, which catalyzes the last step in the carnitine biosynthetic pathway, was lower in *Fmo5^-/-^* mice (Table 3). Although expression of *Cpt1* was similar in *Fmo5^-/-^* and WT mice, that of *Acsl1*, encoding the major hepatic long-chain acyl-CoA synthetase (see above), *Slc25a20* and *Cpt2* was lower in *Fmo5^-/-^* mice. The expression of *Crat*, encoding carnitine acetyl transferase, which catalyzes the reversible transfer of short- and medium-chain acyl groups (C2-C10) between CoA and carnitine in the mitochondrial matrix and regulates the acyl-CoA/CoA ratio, also was lower in *Fmo5^-/-^* mice (Table 3). These results suggest that availability of substrates for β-oxidation was lower in *Fmo5^-/-^* mice.

Expression of *Acadm*, *Acad9* and *Acad11*, encoding acyl-CoA dehydrogenases, which catalyze the first step of β-oxidation, was lower in *Fmo5^-/-^* mice (Table 3). ACADs use electron transfer flavoprotein (ETF) as an electron acceptor to transfer electrons to the mitochondrial electron-transport chain via ETF dehydrogenase. The expression of *Etfdh*, which encodes ETF dehydrogenase was lower in *Fmo5^-/-^* mice (Table 3). Expression of *Hadh*, encoding an enzyme that catalyzes step 3 of β-oxidation, and of *Hadhb* and *Acat1*, which encode enzymes that catalyze step 4, was lower in *Fmo5^-/-^* mice (Table 3).

Fatty acid β-oxidation also takes place in peroxisomes. In these organelles long-chain acyl-CoAs (> C16) are degraded to chain lengths of 6-8 Cs, which are then exported to mitochondria for metabolism to C2 units. Fatty acyl-CoAs are transported into peroxisomes by members of the ATP-binding cassette, sub-family D (ABCD1, 2 and 3). Expression of *Abcd3*, encoding a protein that has a preference for the most hydrophilic fatty acids, was lower in *Fmo5^-/-^*mice (Table 4). The first step in peroxisomal ß-oxidation is catalyzed by acyl-CoA oxidases (ACOXs), steps 2 and 3 by peroxisomal bifunctional enzyme (PBE), encoded by *Hsd17b4*, and step 4 by an acetyl-CoA acyltransferase (ACAA). Expression of *Acox1*, *Hsd17b4* and *Acaa1b* was lower in *Fmo5^-/-^*mice than in WT mice (Table 3). The results suggest that in *Fmo5^-/-^*mice both mitochondrial and peroxisomal β-oxidation are compromised.

**Table 4.**
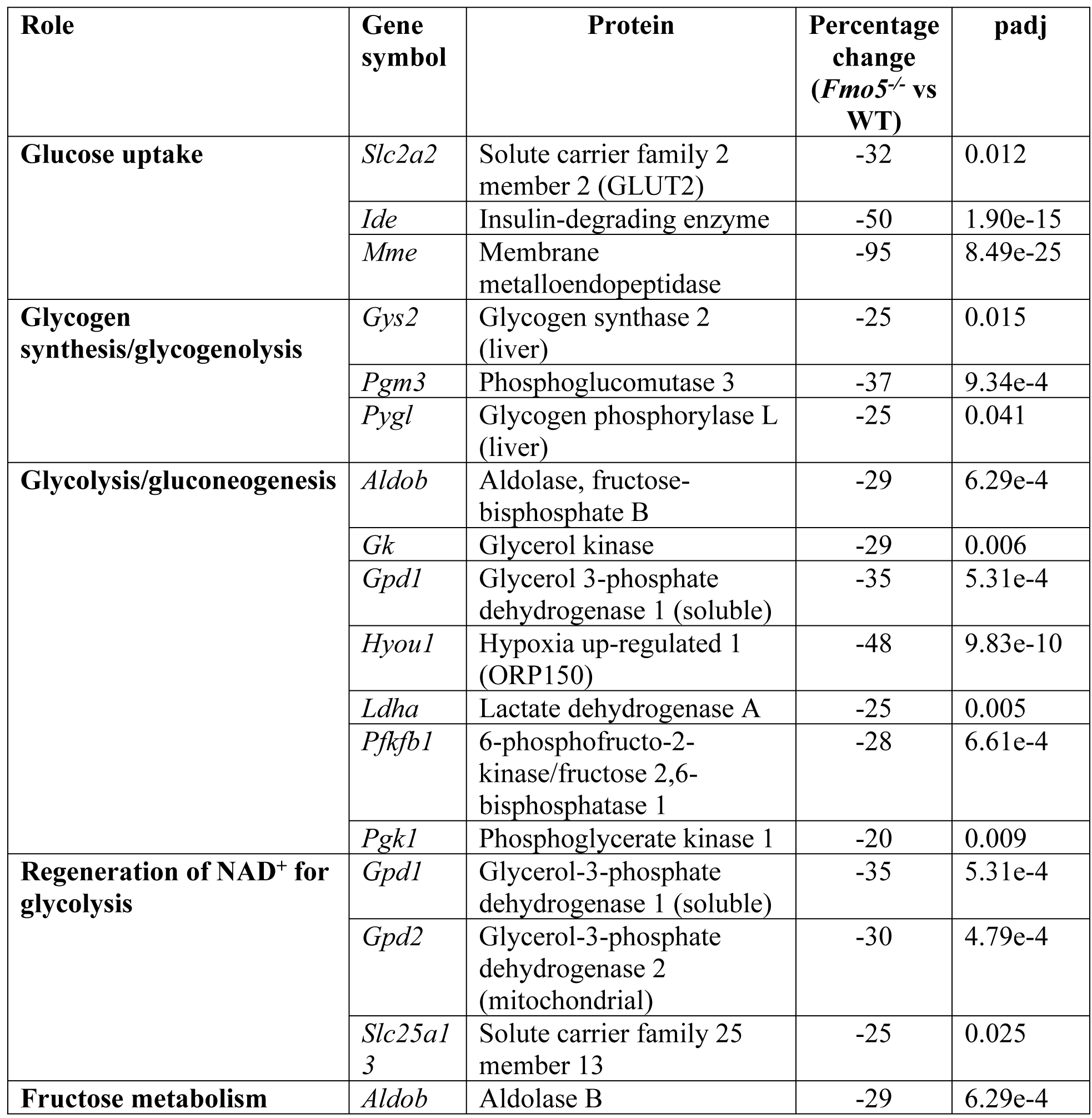
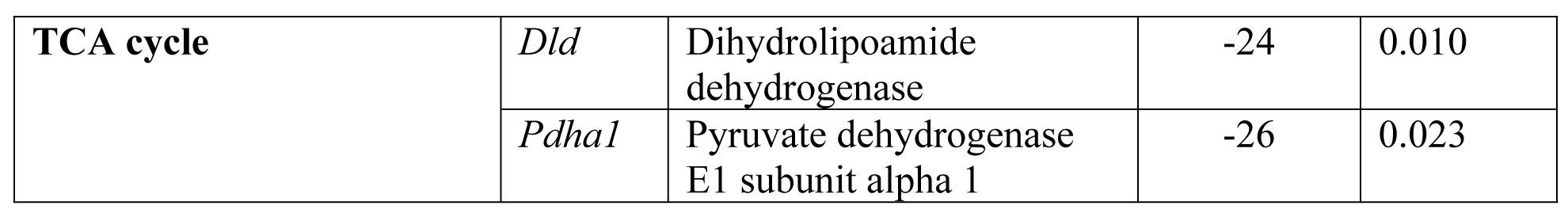
Genes involved in carbohydrate metabolism that are differentially expressed in liver of *Fmo5^-/-^*mice.

### Ketogenesis

Acetyl-CoA generated from mitochondrial fatty acid β-oxidation can be converted to the ketone body acetoacetate via a three-step process. Expression of *Acat1* and *Hmgcs2*, encoding enzymes that catalyze the first two steps, was lower in *Fmo5^-/-^* mice than in WT mice (Table 3), suggesting that *Fmo5^-/-^* mice have a decreased capacity for production of ketone bodies.

### Biosynthesis of TAGs

TAGs are synthesized from glycerol 3-phosphate and fatty acyl-CoAs. The expression of genes encoding three of the enzymes involved in conversion of fatty acids into acyl-CoAs, ACSL1, ACSM3 and ACSS3, was lower in *Fmo5^-/-^*than in WT mice (see Activation of fatty acids to fatty acyl-CoAs, above) (Table 3). Glycerol 3-phosphate is produced from glycerol, via a reaction catalyzed by glycerol kinase (GK), or from dihydroxyacetone phosphate (DHAP), via the action of glycerol 3-phosphate dehydrogenase 1 (GPD1). The expression of genes encoding both GK and GPD1 was lower in *Fmo5^-/-^*mice (Table 3). DHAP is derived from the glycolytic intermediate fructose 1,6-bisphosphate, or from fructose 1-phosphate, by the action of aldolase B, encoded by *Aldob*, the expression of which was lower in *Fmo5^-/-^*mice (Table 3). Fructose 1-phosphate is produced from fructose by the action of ketohexokinase (KHK). Although the abundance of the KHK mRNA was similar in the liver of *Fmo5^-/-^*and WT mice, the abundance of the protein is lower in *Fmo5^-/-^*mice, as is that of aldolase B and GPD1 proteins [12]. The results suggest that hepatic production of the precursors of TAG biosynthesis is compromised in *Fmo5^-/-^*mice. However, although the hepatic concentration of glycerol was lower in *Fmo5^-/-^*mice, that of DHAP and glycerol 3-phosphate was similar in *Fmo5^-/-^*and WT mice (S1 Table).

The main route for TAG biosynthesis is the glycerol 3-phosphate pathway, which consists of four steps. Expression of *Gpam*, which encodes the enzyme that catayzes the first, rate-limiting step, and of *Agpat2* and *Lpin2*, encoding enzymes that catalyze steps 2 and 3, was lower in *Fmo5^-/-^* than in WT mice (Table 3). Expression of *Lpgat1*, encoding lysophosphatidylglycerol acyltransferase 1, which catalyzes formation of diacylglycerol from monoacylglycerol, also was lower in *Fmo5^-/-^* mice (Table 3). The results suggest that biosynthesis of TAGs is compromised in *Fmo5^-/-^* mice.

### TAG storage and secretion

TAGs are stored in lipid droplets. Expression of genes coding for some of the major protein components of lipid droplet coats, perilipin 2, 4 and 5 (*Plin2*, *4* and *5*), was lower in *Fmo5^-/-^*mice (Table 3), suggesting that lipid droplets were less abundant or smaller in these mice. In hepatocytes, lipid droplets are tightly associated with membranes of the ER and components of lipid droplets interact with ER membrane proteins. Expression of *Fitm1*, *Fitm2* and *Cideb*, encoding ER proteins required for lipid droplet formation [24], was lower in *Fmo5^-/-^*mice (Table 3), suggesting that formation of lipid droplets is compromised. *Fitm1* was one of the most significantly differentially expressed genes (padj = 3.71e-10).

A major destination of hepatic TAGs stored in lipid droplets is incorporation into very-low-density lipoprotein (VLDL) particles for secretion, a process that takes place in the ER. Expression of *Copb1*, *Copb2* and *Copg1*, encoding three of five coat complex 1 proteins thought to be involved in transport of TAGs from lipid droplets to the ER, was lower in *Fmo5^-/-^* mice (Table 3). The first step in the process, the addition of TAG to ApoB100 in the ER, to form a primordial VLDL_2_ particle, requires microsomal TAG transfer protein (MTTP), which acts as a heterodimer with prolyl 4-hydroxylase (P4HB). Expression of both *Mttp* and *P4hb* was lower in *Fmo5^-/-^* mice (Table 3). VLDL_2_ particles are converted into VLDL_1_ particles by a second lipidation step, also mediated by MTTP, which involves carboxylesterases (CESs), and expression of *Ces1d*, *Ces1e* and *Ces1f* was lower in *Fmo5^-/-^*mice (Table 3). Expression of *Sar1b*, *Sec23a*, *Sec23b*, *Sec24a* and *Sec24d*, encoding components of coat complex II that interact with CIDEB to transport mature VLDL-TAG particles from the ER to the Golgi, was lower in *Fmo5^-/-^*mice (Table 3), as was that of *Cideb* and *Cd36* (see above), which is also involved in VLDL output from liver [25]. Expression of *Apoa4*, *Apoa5* and *Apoc3* was lower in *Fmo5^-/-^*mice (Table 3). The *Apoa4* was the most significantly differentially expressed gene (padj = 7.00e-27) between *Fmo5^-/-^*and wild-type mice. APOA4 enhances hepatic TAG secretion [26], APOA5 is involved in hepatic TAG accumulation and/or lipid droplet formation [27] and APOC3 promotes assembly and secretion of VLDL particles [28]. The results suggest that production and secretion of VLDL particles is diminished in *Fmo5^-/-^*mice.

### Lipolysis and lipophagy

TAGs in lipid droplets can also be metabolized to fatty acids and glycerol by the sequential action of three enzymes, adipose triglyceride lipase (ATGL), hormone-sensitive lipase and monoglyceride lipase. Expression of *Mgll*, encoding monoglyceride lipase, was lower in *Fmo5^-/-^*than in WT mice (Table 3).

Lipid droplets can be degraded via an autophagic process termed lipophagy. This can be initiated by chaperone-mediated autophagy [29]. The chaperone heat-shock protein 8 (HSPA8) binds to PLIN2 on the coat of lipid droplets and targets them to lysosomes for degradation. Lysosomal-associated membrane protein 2 (LAMP2) is required for fusion of autophagosomes with lysosomes during autophagy. Microtubule associated protein 1 light chain 3α, encoded by the *Map1lc3a* gene, interacts with ATGL and translocates the lipase to the surface of lipid droplets. PNPLA8, a member of the patatin-like phospholipase domain-containing protein family, promotes lipophagy. Expression of *Hspa8*, *Plin2*, *Lamp2*, *Map1lc3a* and *Pnpla8* were all lower in *Fmo5^-/-^*mice (Table 3), as was that of *Lipa*, which encodes lysosomal acid lipase A, the major lipase present in lysosomes. In addition, saturated fatty acids, the concentrations of which were lower in *Fmo5^-/-^*mice (see above and S1 Table and Fig 3B), have been found to activate lipophagy [30]. The results indicate that chaperone-mediated autophagy and lipophagy is less active in *Fmo5^-/-^*mice.

### Cholesterol

The hepatic concentration of cholesterol was higher in *Fmo5^-/-^* mice than in WT mice (S1 Table). Factors that might contribute to this are the lower expression in the liver of *Fmo5^-/-^* mice of genes encoding APOA4, which activates lecithin-cholesterol acyltransferase [31], an enzyme required for cholesterol transport; CD36, which is involved in VLDL output from the liver; bile acid-CoA:amino acid *N*-acyltransferase (*Baat*), which promotes secretion of cholesterol into bile; ABCB4, a P-glycoprotein phosphatidylcholine transporter that is important in bile formation and indirectly determines biliary cholesterol output [32]; and CYP8B1, which is involved in the classic (neutral) bile acid biosynthetic pathway (Table 3). However, despite lower expression of *Cyp8b1* in *Fmo5^-/-^* mice, hepatic concentrations of primary bile acids were similar in *Fmo5^-/-^* and WT mice.

Expression of *Angptl3* was lower in *Fmo5^-/-^*mice (Table 3). ANGPTL3, which is expressed exclusively in liver, is a hepatokine involved in regulating the uptake of lipids, cholesterol and glucose by peripheral tissues [33].

### Glycerophospholipids

The hepatic concentrations of 9 of the 14 phosphatidylethanolamines detected were higher in *Fmo5^-/-^* mice, as were those of phosphoethanolamine, a precursor of phosphatidylethanolamine biosynthesis, and glycerophosphoethanolamine, a breakdown product (S1 Table). Expression of *Apol9a* and *Apol9b*, encoding subunits of APOL9, which binds phophatidylethanolamines [34], was higher in *Fmo5^-/-^* mice (Table 3).

Phosphatidylethanolamines, along with serine, the concentration of which was also elevated in *Fmo5^-/-^* mice, are precursors in the biosynthesis of phosphatidylserines, and the concentrations of two of three phosphatidylserines detected were also elevated in these mice (S1 Table). Of 27 lysophospholipids detected, the concentrations of three were higher and nine lower in *Fmo5^-/-^* mice. However, differences were consistent only with lysophosphatidylcholines, with six of the nine detected being lower in *Fmo5^-/-^* mice (S1 Table).

The hepatic concentrations of six of the eight plasmalogens detected were higher in *Fmo5^-/-^* mice (S1 Table). Plasmalogens are synthesized in the liver and transported to other tissues by lipoproteins. The predicted decrease in production of VLDL particles (see above) would negatively affect export of plasmalogens from the liver.

### Sphingolipids

The hepatic concentration of serine, a precursor of sphingolipid biosynthesis, and of two intermediates in ceramide biosynthesis, sphinganine (dihydrosphingosine) and *N*-palmitoyl-sphinganine (d18:0/16:0) (a major dihydroceramide), was higher in *Fmo5^-/-^* mice (S1 Table and Fig 3D). The concentrations of dihydrosphingomyelins, which can be produced from dihydrosphingosine, were also higher in *Fmo5^-/-^*mice, by as much as 22-fold (S1 Table and Fig 3D). Differences were also observed between *Fmo5^-/-^*and WT mice in the hepatic concentrations of several ceramides, hexosylceramides and sphingomyelins (S1 Table).

The concentration of sphingosine, a catabolic product of ceramides, was higher in *Fmo5^-/-^* mice (S1 Table). The breakdown of sphingolipids proceeds via the conversion of sphingosine to sphingosine 1-phosphate, which is then converted to phosphoethanolamine and fatty aldehydes. The lower expression in *Fmo5^-/-^* mice of *Aldh3a2*, *Acsl1* and *Slc27a2* (Table 3), encoding aldehyde dehydrogenase 3A2, acyl-CoA synthetase long-chain 1 and very long-chain acyl-CoA synthetase, three of the enzymes involved in the subsequent metabolism of fatty aldehydes to fatty acyl-CoAs, might contribute to the higher hepatic concentration of sphingosine in these animals.

### Eicosanoids

The hepatic concentration of the omega-6 polyunsaturated fatty acid arachidonate, a precursor of pro-inflammatory eicosanoids, was lower in *Fmo5^-/-^*mice (S1 Table), as was expression of *Pla2g6* and *Pnpla8*, which encode enzymes involved in the selective release of arachidonate from glycerophospholipids [35], and *Ephx1* and *Abhd6*, encoding enzymes that respectively metabolize 2- and 1-arachidonylglycerol to arachidonate and glycerol, both of which were present at lower concentrations in *Fmo5^-/-^*mice (Table 3). Expression of *Ptgs1*, encoding prostaglandin synthase 1, which catalyzes the first step in the production of prostaglandins from arachidonate, was higher in *Fmo5^-/-^* mice (Table 3), as were the hepatic concentrations of two metabolites of arachidonate, 12-hydroxyeicosatetraenoate (12-HETE) and 12-hydroxyheptadecatrienoate (12-HHTrE) (S1 Table), indicating that decreased production and increased metabolism contribute to the lower arachidonate concentration in *Fmo5^-/-^* mice.

### Non-lipid metabolism

Metabolomic and transcriptomic data also revealed differences between *Fmo5^-/-^* and WT mice in the hepatic concentrations of several non-lipid metabolites (S1 Table and Fig 4A) and in the expression of genes encoding proteins involved in non-lipid metabolic pathways.

### Glucose uptake from plasma

The concentration of glucose in both the liver and plasma (S1 and S2 Tables) was lower in *Fmo5^-/-^*mice than in WT mice. Expression of *Slc2a2*, encoding the glucose transporter GLUT2, which because of its low affinity for glucose is considered a glucose sensor, was lower in *Fmo5^-/-^*mice, which is consistent with the lower intrahepatic glucose concentration of these mice.

Among the genes that exhibited the most significantly lower expression in *Fmo5^-/-^* mice were *Ide* (padj = 1.90e-15), encoding insulin-degrading enzyme, a metalloendopeptidase whose substrates include insulin and glucagon [36], and *Mme* (padj = 8.49e-25), encoding a cell-surface metalloendopeptidase, whose substrates include glucagon [37] (Table 4). The lower expression of *Ide* in *Fmo5^-/-^*mice might be in response to the lower plasma concentration of insulin. Despite the consequential potential decrease in glucagon degradation, *Fmo5^-/-^*mice have lower plasma and liver concentrations of glucose.

### Glycogen synthesis/glycogenolysis

Expression of *Gys2*, encoding hepatic glycogen synthase, which catalyzes the rate-limiting step in glycogen synthesis, *Pygl*, encoding liver glycogen phosphorylase, which catalyzes the release of glucose 1-phosphate from glycogen, and *Pgm3*, encoding phosphoglucomutase 3, which catalyzes the interconversion of glucose 1-phosphate and glucose 6-phosphate and is thus involved in both glycogen formation and utilization, was lower in *Fmo5^-/-^* mice (Table 4), suggesting that disruption of the *Fmo5* gene perturbs glycogen metabolism.

### Glycolysis/Gluconeogenesis

Expression of *Aldob* and *Pgk1* was lower in *Fmo5^-/-^*mice (Table 4). The genes encode aldolase B and phosphoglycerate kinase 1, which respectively catalyze the interconversion of fructose 1,6-bisphosphate and glyceraldehyde 3-phosphate plus DHAP, and of 1,3-bisphoglycerate and 3-phosphoglycerate, reversible reactions in glycolysis and gluconeogenesis.

A key point of control in the glycolytic and gluconeogenic pathways is the reciprocal regulation of phosphofructokinase, which catalyzes the conversion of fructose 6-phosphate to fructose 1,6-bisphosphate, an irreversible step that commits glucose to glycolysis, and fructose 1,6-bisphosphatase, which catalyzes the reverse reaction in gluconeogenesis.

Phosphofructokinase is allosterically inhibited by citrate and activated by ADP, AMP and fructose 2,6-bisphosphate, whereas fructose 1,6-bisphosphatase is activated by citrate and inhibited by ADP, AMP and fructose 2,6-bisphosphate. The hepatic concentration of citrate was higher and that of ADP was lower in *Fmo5^-/-^* mice (S1 Table), suggesting that in *Fmo5^-/-^* mice glycolysis would be suppressed and gluconeogenesis stimulated. This is supported by the lower expression in *Fmo5^-/-^* mice of *Hyou1* (Table 4), one of the most significantly differentially expressed genes (padj = 9.83e-10), encoding ORP150, which has an inhibitory effect on gluconeogenesis.

Fructose 2,6-bisphosphate is produced via a reaction catalyzed by the bifunctional enzyme 6-phosphofructo-2-kinase/fructose 2,6-biphosphatase 1, encoded by *Pfkfb1*. The kinase activity of the enzyme is inhibited by phosphorylation catalyzed by cAMP-dependent protein kinase A, encoded by *Prkaca*. The enzyme is also inhibited by itaconate, the concentration of which was 10-fold higher in *Fmo5^-/-^* mice (S1 Table). Expression of *Pfkfb1* was lower in *Fmo5^-/-^* mice, but that of *Prkaca* was similar in *Fmo5^-/-^* and WT mice. The data suggest that production of fructose 2,6-bisphosphate would be lower in *Fmo5^-/-^* mice than in WT mice, leading to suppression of glycolysis and stimulation of gluconeogenesis. The latter might be in response to the lower plasma and hepatic glucose concentrations in *Fmo5^-/-^* mice (S1 and S2 Tables).

Glycerol can be converted into DHAP, by reactions catalyzed by glycerol kinase and glycerol 3-phosphate dehydrogenase 1, encoded by *Gk* and *Gpd1* respectively, and is thus a substrate for both glycolysis and gluconeogenesis. The hepatic concentration of glycerol and the expression of *Gk* and *Gpd1* was lower in *Fmo5^-/-^*mice (see TAG biosynthesis above and Table 4). Lactate can be converted into pyruvate, by the action of lactate dehydrogenase, encoded by *Ldha*, and is thus a substrate for gluconeogenesis. The hepatic concentration of lactate and the expression of *Ldha* was lower in *Fmo5^-/-^*mice (S1 Table and Table 4). The NAD^+^ utilized by glyceraldehyde 3-phosphate dehydrogenase during glycolysis can be regenerated via the glycerophosphate and malate-aspartate shuttles. In the glycerophosphate shuttle, NAD^+^ is produced from NADH during the reduction of DHAP to glycerol 3-phosphate, catalyzed by cytosolic glycerol 3-phosphate dehydrogenase, encoded by *Gpd1*. Glycerol 3-phosphate is converted back to DHAP by mitochondrial glycerol 3-phosphate dehydrogenase, encoded by *Gpd2*, in the process reducing FAD to FADH_2_. Expression of *Gpd1* and *Gpd2* was lower in *Fmo5^-/-^* mice (Table 4), as was the concentration of FAD (S1 Table). Expression of *Slc25a13*, which encodes a component of the malate-aspartate shuttle that translocates aspartate from mitochondria to cytosol, was lower in *Fmo5^-/-^*mice (Table 4). The results suggest that in *Fmo5^-/-^*mice activity of both shuttles is decreased, which would compromise regeneration of NAD^+^and, thus, contribute to suppression of glycolysis.

Of the 18 glucogenic amino acids, the hepatic concentrations of 13 were higher in *Fmo5^-/-^* mice (S1 Table), supporting the suggestion that gluconeogenesis is stimulated in these mice. In addition, the concentrations of fumarate and malate, which can be converted into oxaloacetate and, thus, are precursors of gluconeogenesis, were higher in *Fmo5^-/-^* mice (S1 Table).

### Fructose metabolism

Fructose is metabolized to fructose 1-phosphate by a reaction catalyzed by fructokinase, encoded by *Khk*, and subsequently to glyceraldehyde and DHAP, catalyzed by aldolase B, encoded by *Aldob*. Although expression of *Khk* was similar in *Fmo5^-/-^* and WT mice, the protein is less abundant in *Fmo5^-/-^* mice [12]. Expression of *Aldob* was lower in *Fmo5^-/-^* mice (see above and Table 4), as was the abundance of the protein [12]. The data suggest that fructose metabolism is compromised in *Fmo5^-/-^* mice.

### Tricarboxylic acid (TCA) cycle

In comparison with WT mice, *Fmo5^-/-^* mice had higher hepatic concentrations of the TCA cycle intermediates citrate, aconitate, fumarate and malate (S1 Table), suggesting elevated utilization of the cycle. *Fmo5^-/-^* mice also had a higher concentration of itaconate (∼10-fold) (see Glycolysis/Gluconeogenesis above) (S1 Table), derived from the TCA cycle intermediate *cis*-aconitate. Expression of *Pdha1* and *Dld*, which encode two of the five components of the pyruvate dehydrogenase complex, was lower in *Fmo5^-/-^* mice (Table 4), suggesting that elevated TCA cycle intermediates would be derived from sources other than pyruvate. Although itaconate inhibits the activity of succinate dehydrogenase, the concentration of the product of the reaction catalyzed by this enzyme, fumarate, was higher in *Fmo5^-/-^* mice, as was that of malate (S1 Table), the product of the subsequent reaction of the cycle, possibly as a consequence of increased flow through the aspartate-arginosuccinate shunt [38], suggested by the higher concentrations in *Fmo5^-/-^* mice of aspartate and arginosuccinate (S1 Table). Other potential sources of TCA cycle intermediates are amino acids, many of which are present in higher concentrations in *Fmo5^-/-^* mice (S1 Table).

### Amino acid metabolism

The hepatic concentrations of 15 of 20 amino acids were higher in *Fmo5^-/-^*than in WT mice (S1 Table). This may be a consequence of the greater consumption of food by *Fmo5^-/-^*mice [12], but decreased degradation may also contribute. The expression of *Bckdha*, encoding the alpha subunit of branched chain alpha-keto acid dehydrogenase E1, which catalyzes the second, rate-limiting step in the degradation of isoleucine, valine and leucine, was lower in *Fmo5^-/-^* mice, as was that of *Hibch* and *Hibadh*, encoding enzymes of the valine degradation pathway, *Acat1* and *Mccc1*, encoding enzymes of the isoleucine and leucine degradative pathways, respectively, and *Tdo2*, encoding the first, rate-limiting step in tryptophan degradation (Table 5).

**Table 5.** Genes encoding proteins involved in amino acid metabolism that are differentially expressed in *Fmo5^-/-^* mice. NAD^+^.

| Gene symbol | Protein | Percentage change (*Fmo5*<sup>-/-</sup> vs WT) | padj |
| --- | --- | --- | --- |
| *Acat1* | Acetyl-CoA acetyltransferase 1 | -28 | 2.20e-4 |
| *Arg1* | Arginase 1 | -21 | 0.037 |
| *Asl* | Argininosuccinate lyase | -22 | 0.039 |
| *Bckdha* | Branched chain keto acid dehydrogenase E1 alpha | -22 | 0.045 |
| *Hibadh* | 3-hydroxybutyrate dehydrogenase | -19 | 0.049 |
| *Hibch* | 3-Hydroxyisobutyryl-CoA hydrolase | -21 | 0.037 |
| *Mccc1* | Methylcrotonyl -CoA carboxylase subunit 1 | -39 | 1.22e-8 |
| *Tdo2* | Tryptophan 2, 3 dioxygenase | -29 | 0.002 |

The hepatic concentrations of two components of the urea cycle, ornithine and aspartate, and of two products of the cycle, urea and fumarate, were higher in *Fmo5^-/-^* mice (S1 Table). However, expression of *Asl* and *Arg1*, encoding arginosuccinate lyase and arginase 1, which catalyze the formation of fumarate and of urea and ornithine, respectively, was lower in *Fmo5^-/-^* mice (Table 5).

Although the hepatic concentration of quinolinate, a major precursor of NAD^+^ and NADP^+^ biosynthesis, was lower in *Fmo5^-/-^* than in WT mice (S1 Table and Fig 4B), that of NAD^+^ was similar in *Fmo5^-/-^* and WT mice (S1 Table). The hepatic concentration of nicotinamide, which is both a major breakdown product of NAD^+^, and, via the salvage pathway, a precursor of NAD^+^ [39], was lower in *Fmo5^-/-^* mice (S1 Table and Fig 4B). Nicotinamide can be further metabolized, via the NAD^+^ clearance pathway, to 1-methylnicotinamide and, subsequently, to pyridones by a reaction catalyzed by AOX1 [39]. The hepatic concentration of 1-methylnicotinamide was higher in *Fmo5^-/-^* mice (S1 Table and Fig 4B). Expression of *Aox1* was lower in *Fmo5^-/-^* mice (Table 1), which would contribute to the accumulation of 1-methylnicotinamide in these animals. Nicotinamide can also be converted to its *N*-oxide, by a reaction catalyzed by CYP2E1 [40]. However, the decreased ratio of nicotinamide *N*-oxide to 1-methylnicotinamide in *Fmo5^-/-^* mice, together with the similar level of *Cyp2e1* expression in *Fmo5^-/-^* and wild-type mice, suggests that FMO5 plays a role in catalyzing the *N*-oxygenation of nicotinamide.

### ADP-ribose and ADP-ribosylation

One of the uses of NAD^+^ is as a cofactor for the ADP-ribosylation of proteins. The hepatic concentration of ADP-ribose was 95% lower in *Fmo5^-/-^* mice (S1 Table and Fig 4B), the largest decrease of any metabolite measured. Although expression of genes encoding proteins involved in removal of ADP-ribose from proteins, *Adprh*, *MacroD1*, *MacroD2*, *Oard1 A* and *Parg* [41], was similar in *Fmo5^-/-^* and WT mice, that of *Parp10* and *Parp14*, encoding ARTD10 and ARTD8, which catalyze mono-ADP ribosylation (MARylation) of proteins [42], was higher in *Fmo5^-/-^* mice (Table 6), suggesting that the substantially lower concentration of ADP-ribose in these animals could be a consequence of increased MARylation of proteins.

**Table 6.**
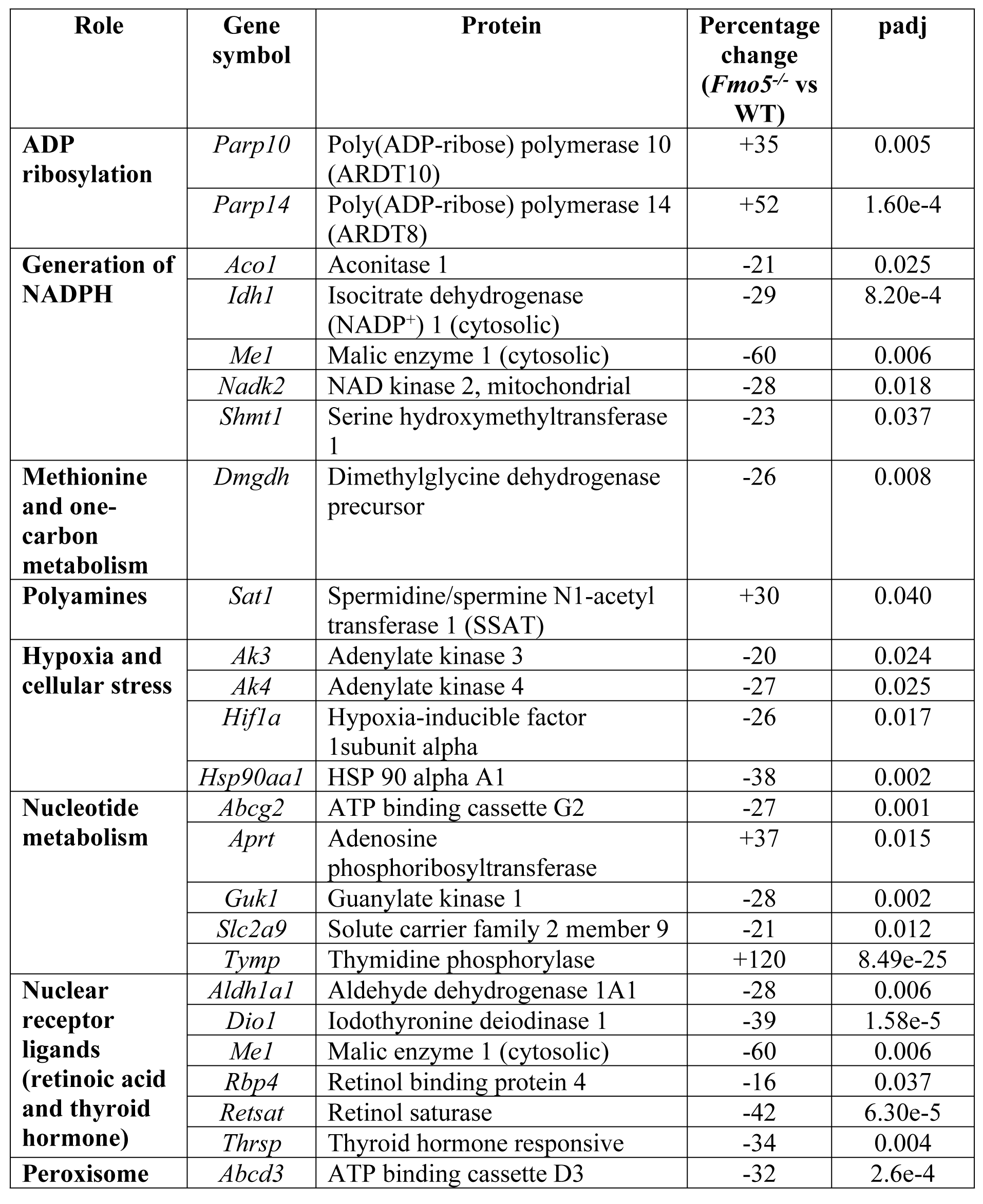
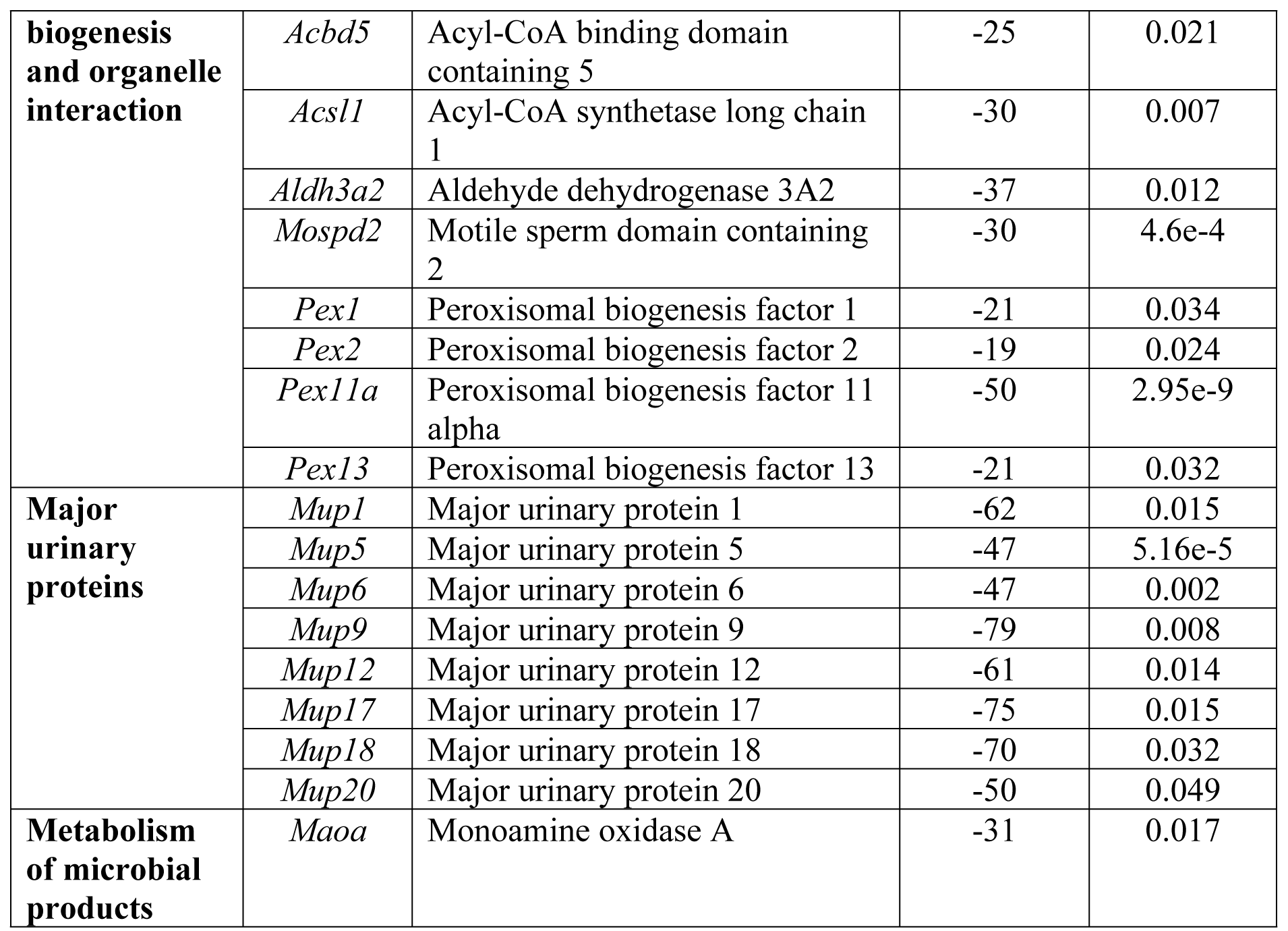
Genes encoding proteins involved in other metabolic processes that are differentially expressed in *Fmo5^-/-^*mice.

### Generation of NADPH

Expression of mitochondrial NAD kinase, encoded by *Nadk2*, which converts NAD^+^ to NADP^+^, was lower in *Fmo5^-/-^* mice (Table 6), suggesting that, although the concentration of NAD^+^ was similar in *Fmo5^-/-^* and WT mice (S1 Table), production of NADP^+^ in mitochondria is compromised in *Fmo5^-/-^* mice.

NADPH can be generated from NADP^+^ by several means. The pentose phosphate pathway is a major source of cytoplasmic NADPH. NADPH can also be generated from NADP^+^ in the cytosol or mitochondria by conversion of malate to pyruvate, catalyzed by malic enzyme (ME), conversion of isocitrate to α-ketoglutarate, catalyzed by isocitrate dehydrogenase (IDH), or during one-carbon metabolism. Expression of *Me1* and *Idh1*, encoding the cytosolic forms of ME and IDH, was lower in *Fmo5^-/-^* mice, as was that of *Aco1*, encoding aconitase 1 (Table 6), which catalyzes production of isocitrate from citrate in the cytoplasm. Expression of *Idh2*, encoding mitochondrial IDH, was similar in *Fmo5^-/-^* and WT mice, whereas that of *Me3*, encoding mitochondrial ME, was below the limit of detection.

Generation of NADPH via one-carbon metabolism occurs in both the cytosol and mitochondria during conversion of 5,10-methylene tetrahydrofolate to 5,10-methenyl tetrahydrofolate, and in mitochondria also during conversion of 10-formyl tetrahydrofolate to tetrahydrofolate. To form 5,10-methylene tetrahydrofolate, a carbon unit is transferred from glycine or serine in a reaction catalyzed by serine hydroxymethyltransferase. Expression of *Shmt1*, encoding the cytosolic form of the enzyme, but not that of *Shmt2*, encoding the mitochondrial form, was lower in *Fmo5^-/-^* mice (Table 6). The results suggest that generation of cytosolic NADPH via ME, IDH and one-carbon metabolism is suppressed in *Fmo5^-/-^* mice.

### Methionine and one-carbon metabolism

Methionine is involved in one-carbon metabolism, via its conversion to the methyl donor *S*-adenosylmethionine (SAM) and interaction of the methionine cycle with the folate cycle (Fig 5). The methionine cycle is also linked to the homocysteine transsulfuration pathway, producing cysteine, and the methionine salvage cycle, which intersects with polyamine biosynthesis, via decarboxylation of ornithine to produce putrescine, a precursor of spermidine and spermine (Fig 5). Differences were observed between *Fmo5^-/-^*and WT mice in the hepatic concentrations of several one-carbon metabolism intermediates: those of methionine and *N*^5^-methyl-tetrahydrofolate, a precursor of methionine synthesis via homocysteine, were higher in *Fmo5^-/-^*mice, whereas folate and 7,8-dihydrofolate, precursors of tetrahydrofolate, were lower in these animals, as were sarcosine and dimethylglycine (S1 Table), and expression of *Dmgdh*, which encodes the enzyme that catalyzes conversion of dimethylglycine to sarcosine, was lower in *Fmo5^-/-^* mice. *Fmo5^-/-^* mice also had higher hepatic concentrations of guanidinoacetate, creatine and cysteine (S1 Table and Fig 5). Methionine and creatine also function as antioxidants [43,44]. Methionine interacts with ROS to produce methionine sulfoxide [43], the hepatic concentration of which was higher in *Fmo5^-/-^* mice (S1 Table and Fig 5). Collectively, these results indicate that in *Fmo5^-/-^* mice the metabolism of 1-carbon units is altered, and intermediates appear to be channelled towards creatine and *N*^5^-methyl-tetrahydrofolate.

**Fig 4.**
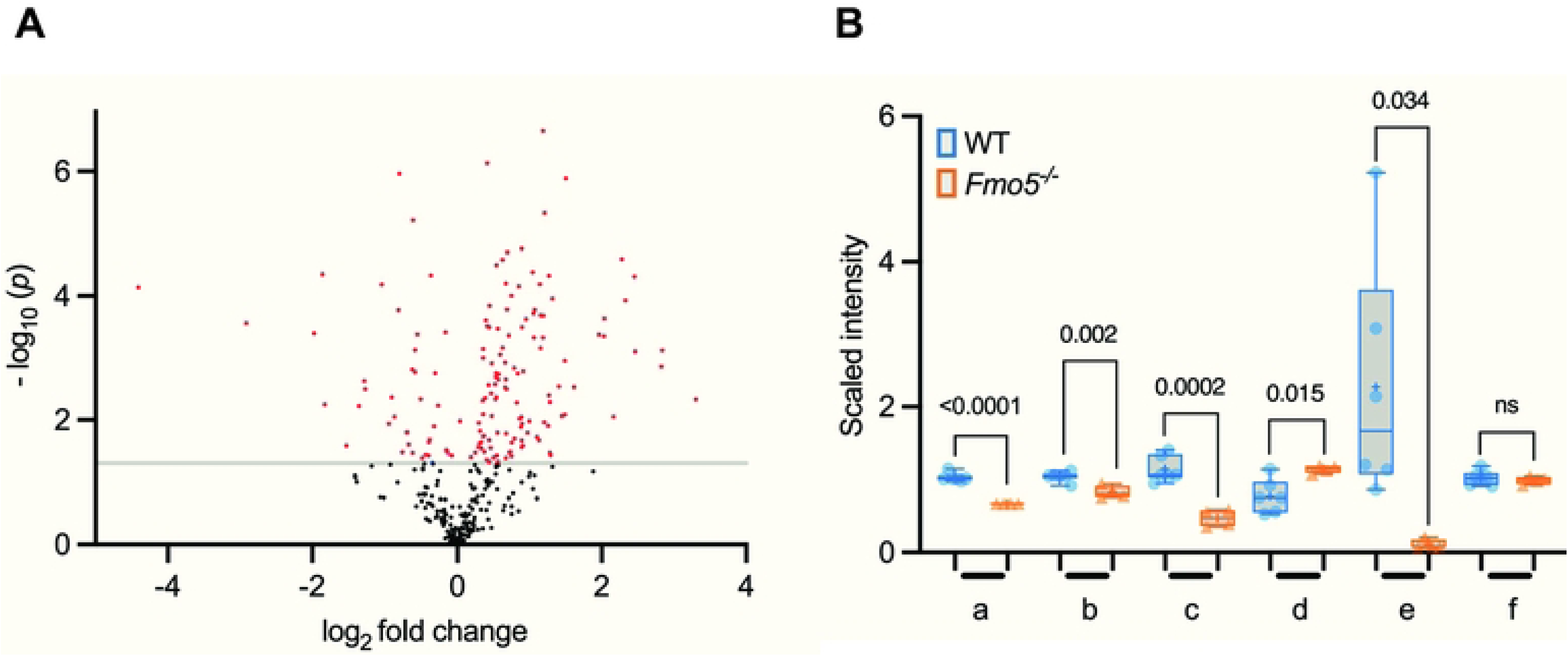
Differences in abundance of non-lipid molecules between *Fmo5^-/-^* vs WT mice. (A) Volcano plot representation of non-lipid molecules (log_2_-fold change) of *Fmo5^-/-^* vs WT mice. Red dots are non-lipids that are significantly increased or decreased in *Fmo5^-/-^* mice with a -log_10_ (*p*) above 1.301. (B) Nicotinamide and associated metabolites. a. quinolinate; b. nicotinamide; c. nicotinamide *N*-oxide; d. 1-methylnicotinamide; e. adenosine 5’-diphosphoribose (ADP-ribose); f. nicotinamide adenine dinucleotide (NAD+). + indicates mean value, numbers above box plot are *p* values. ns = not significant

**Fig 5.**
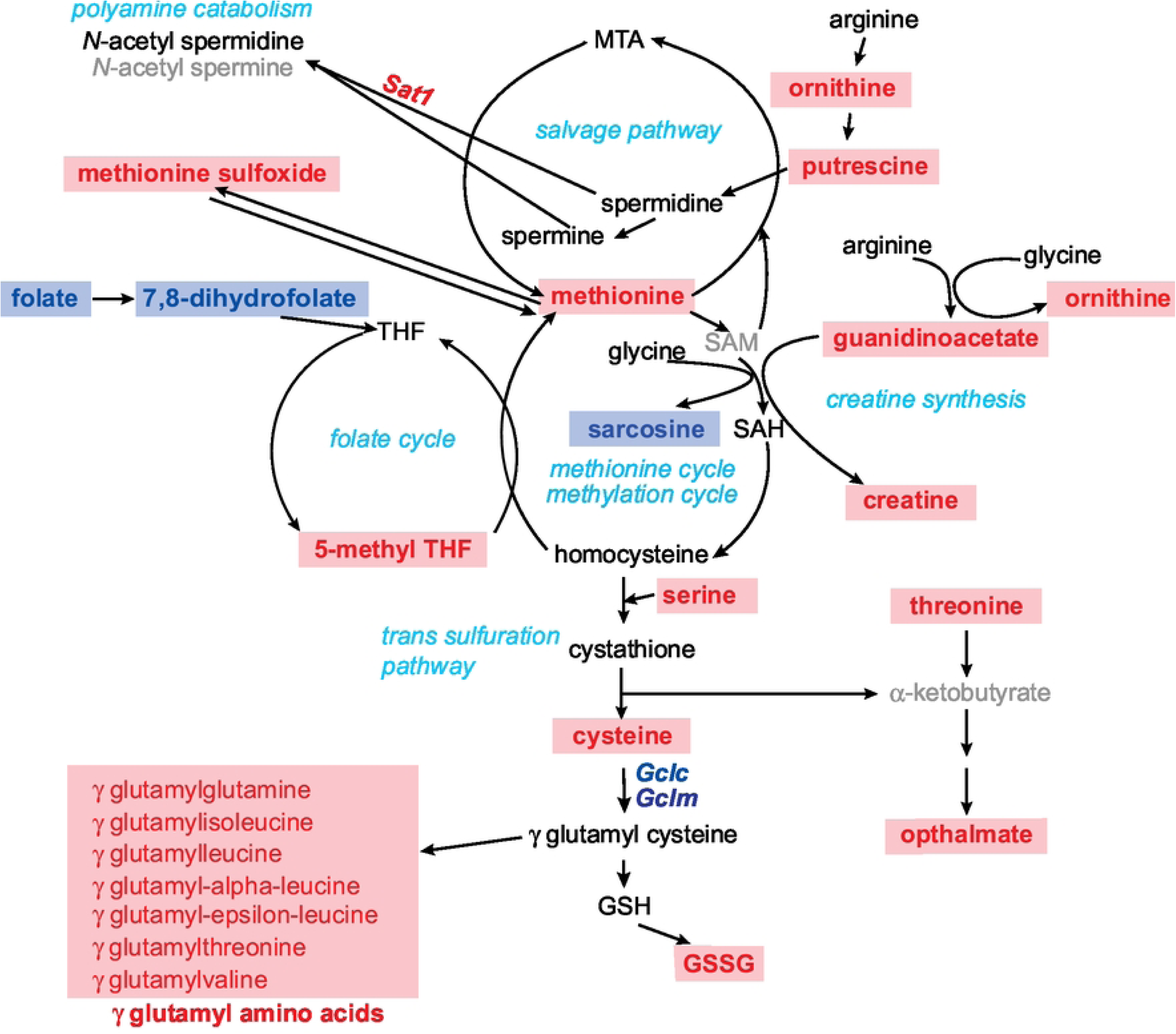
One-carbon metabolism perturbations in *Fmo5^-/-^* mice.

Red indicates an increase, blue a decrease and black no change in metabolite abundance or gene expression in *Fmo5^-/-^*vs WT mouse liver. Grey indicates not measured. Gene symbols are in italics. Pathways/processes are named in turquoise. GSH, glutathione; GSSG, oxidized glutathione; MTA, 5’-methyl-thioadenosine; SAH, S-adenosylhomocysteine; SAM, S-adenosylmethionine; THF, tetrahydrofolate.

### Glutathione

The hepatic concentration of reduced glutathione (GSH) was similar in *Fmo5^-/-^* and WT mice, but that of oxidized glutathione (GSSG) was higher in *Fmo5^-/-^* mice (S1 Table and Fig 5). The concentration of cysteine, one of the precursors of glutathione, was higher in *Fmo5^-/-^* mice, but expression of *Gclc* and *Gclm*, encoding the two components of glutamate-cysteine ligase, which catalyzes the first, rate-limiting, step in glutathione synthesis, were both lower in *Fmo5^-/-^* mice (Table 1). In addition, the hepatic concentration of ophthalmate, an analogue of glutathione, which acts as an anti-oxidant, was higher in *Fmo5^-/-^* mice (S1 Table and Fig 5). Hepatic concentrations of 7 of the 11 detected γ-glutamyl amino acids, which are formed by transfer of a glutamyl group from GSH, were higher in *Fmo5^-/-^* mice (S1 Table and Fig 5).

### Polyamines

Hepatic concentrations of the polyamine putrescine and its precursor ornithine were higher in *Fmo5^-/-^* mice (S1 Table and Fig 5), as was expression of *Sat1* (Table 6 and Fig 5), which encodes SSAT, an acetyltransferase that catalyzes the rate-limiting step in polyamine catabolism, producing acetylated derivatives of spermine and spermidine, which are converted back to putrescine [45]. Enhanced expression of *Sat1* could account for the increased concentration of putrescine in *Fmo5^-/-^* mice.

### Hypoxia and cellular stress

SSAT decreases the abundance of hypoxia-inducible factor 1A (HIF1A) [46]. Although this occurs via SSAT binding to HIF1A and promoting its ubiquitination and degradation, it is of interest that the abundance of HIF1A mRNA was lower in *Fmo5^-/-^*mice (Table 6). Degradation of HIF1A protein is also induced by inhibition of HSP90 [46], and expression of *Hsp90aa1* was lower in *Fmo5^-/-^* mice (Table 6). Thus, increased expression of *Sat1* and decreased expression of *Hsp90aa1* and *Hif1a* would contribute to a decrease in abundance of HIF1A in *Fmo5^-/-^* mice. HIF1A upregulates transcription of *Pgk1* and *Ldha* [47]. Hence, the predicted lower abundance of HIF1A in *Fmo5^-/-^* mice might account for the lower expression of *Pgk1* and *Ldha* in these animals (see Glycolysis/Gluconeogenesis above and Table 4).

Two other genes induced by HIF1A are *Ak3* and *Ak4* [48,49], encoding mitochondrial matrix adenylate kinases. Expression of both *Ak3* and *Ak4* was lower in *Fmo5^-/-^*mice (Table 6). Knockdown of *Ak4* increases concentrations of the TCA cycle intermediates fumarate and malate [49]. Thus, lower expression of *Ak4* might account for the higher hepatic concentrations of fumarate and malate in *Fmo5^-/-^* mice (S1 Table).

### Nucleotide metabolism

The enzymes encoded by *Ak3* and *Ak4* catalyze the production of ADP from AMP. Thus, the lower expression of the genes in *Fmo5^-/-^*mice (see Hypoxia and cellular stress above and Table 6) might account for the lower hepatic concentration of ADP in these animals (see Glycolysis/Gluconeogenesis above and S1 Table). Similarly, expression of *Guk1*, which encodes the enzyme that catalyzes production of GDP from GMP, was lower in *Fmo5^-/-^*mice (Table 6).

AMP is broken down to hypoxanthine, then xanthine, which is also a breakdown product of GMP. Xanthine is metabolized to urate and then, in mouse, to allantoin. Hepatic concentrations of hypoxanthine, xanthine, urate and allantoin were all higher in *Fmo5^-/-^* mice (S1 Table), indicating that mice lacking FMO5 have an increased catabolism of purines. The higher expression in *Fmo5^-/-^* mice of *Aprt* (Table 6), encoding adenosine phosphoribosyltransferase, which converts adenine to AMP in a purine salvage pathway, might be an attempt to restore AMP. Lower expression in *Fmo5^-/-^* mice of *Abcg2* and *Slc2a9* (Table 6), which encode urate transporters, might contribute to the higher hepatic urate concentration of these animals.

The hepatic concentration of orotate, a precursor of pyrimidine ring synthesis, was higher in *Fmo5^-/-^* mice, but concentrations of CMP and UTP were not elevated in these animals (S1 Table). The hepatic concentrations of cytidine, cytosine, uracil and thymine, initial breakdown products of pyrimidine nucleotides, were higher in *Fmo5^-/-^* mice (S1 Table), as was expression of *Tymp* (Table 6), encoding thymidine phosphorylase, which catalyzes the catabolism of thymidine to thymine [50]. *Tymp* was one of the most significantly differentially expressed genes (padj = 8.49e-25). The results suggest an increased rate of pyrimidine nucleotide metabolism in these animals. However, the concentration of 3-ureidopropionate, a later breakdown product of cytosine and uracil, was lower in *Fmo5^-/-^* mice.

### Nuclear receptor ligands (retinoic acid and thyroid hormone)

Expression of several genes encoding proteins involved in the transport or metabolism of retinol and thyroxine was lower in *Fmo5^-/-^*mice (Table 6): *Rbp4*, encoding retinoic acid-binding protein-4 (RBP4), which is synthesized mainly in liver and transports retinol in the plasma; *Retsat*, which encodes retinol saturase, an ER oxidoreductase that catalyzes the conversion of retinol to 13,14-dihydroretinol; and *Aldh1a1*, encoding aldehyde dehydrogenase 1A1, which catalyzes the conversion of all-trans retinal to all-trans retinoic acid, the rate-limiting step in production of the main endogenous active retinoic acid metabolite. The lower expression of *Aldh1a1* might contribute to the higher hepatic concentration of retinal in *Fmo5^-/-^* mice (S1 Table 1).

The expression of *Dio1*, encoding iodothyronine deiodinase 1, which catalyzes the hepatic conversion of thyroxine (T4) to triiodothyronine (T3), providing a source of active thyroid hormone for the plasma, also was lower in *Fmo5^-/-^* mice, as was that of the thyroid hormone-inducible genes *Me1* (see above) and *Thrsp* (Table 6), which encodes thyroid hormone responsive protein.

### Peroxisome biogenesis and organelle interaction

Expression of *Pex1*, *Pex2*, *Pex11a* and *Pex13*, encoding factors involved in biogenesis of and import of proteins into peroxisomes, was lower in *Fmo5^-/-^* mice (Table 6), suggesting that disruption of the *Fmo5* gene may compromise the formation and maturation of peroxisomes. *Pex11a* was one of the most significant differentially expressed genes (padj = 2.95e-9).

Expression of *Acbd5*, which encodes acyl-CoA binding domain-containing protein 5 (ACBD5), one of the main proteins involved in tethering peroxisomes to the ER and in the motility and positioning of peroxisomes [51], was lower in *Fmo5^-/-^* mice (Table 6), as was that of *Mospd2*, which encodes an ER protein thought to be involved in tethering to peroxisomes via interaction with ACBD5 [52]. Another ER protein that interacts with ACBD5 is acyl-CoA synthetase long-chain family member 1 (ACSL1), which also interacts with fatty aldehyde dehydrogenase (encoded by *Aldh3a2*), present in both the ER and peroxisomes, and ABCD3, the peroxisomal branched-chain fatty acid transporter. Expression of *Acsl1*, *Aldh3a2* and *Abcd3* was lower in *Fmo5^-/-^* mice (Table 6), suggesting that channelling of fatty acids between the ER and peroxisomes may be compromised in these animals.

### Major urinary proteins (MUPs)

MUPs are synthesized mainly in the liver, and in C57BL6 mice comprise 3.5-4% of total protein synthesis in male liver [53]. Of 14 *Mup* genes, the expression of eight was lower in *Fmo5^-/-^*mice (Table 6). MUPs function as transport proteins and pheromones; they promote aggression in male mice and the lower expression of these genes may account for the more placid behaviour of *Fmo5^-/-^*mice that we have observed. MUP20 is sexually attractive to females. Although the expression of *Mup20* was lower in *Fmo5^-/-^*mice, the knockouts displayed no problems with breeding.

### Microbial products and food components

Several products of gut microbial metabolism of tyrosine and tryptophan that have subsequently undergone host conjugation reactions by sulfation (p-cresyl sulfate, phenol sulfate and 3-indoxyl sulfate) or glucuronidation (p-cresol glucuronide and phenol glucuronide) were present in higher concentrations in the livers of *Fmo5^-/-^* mice (S1 Table 1 and Fig 6). However, none of the transcripts of genes encoding UDP glucuronosyl transferases (*Ugt*) or sulfotransferases (*Sult*) was more abundant in the livers of *Fmo5^-/-^* mice. The hepatic concentration of serotonin, a product of host and microbial metabolism of tryptophan [54], was also higher in *Fmo5^-/-^* mice than in WT mice (S1 Table) and expression of *Maoa*, encoding monoamine oxidase-A, which catalyzes the metabolism of serotonin, was lower in *Fmo5^-/-^* mice. The hepatic concentration of hydroxyphenyllactate, also a product of microbial tyrosine metabolism, was lower in *Fmo5^-/-^* mice (S1 Table). Of the 19 metabolites categorised as food components, the hepatic concentrations of seven were lower and one was higher in *Fmo5^-/-^* mice (S1 Table). Differences between *Fmo5^-/-^* and WT mice in the hepatic abundance of host-modified microbial products might be a consequence of differences in composition of the gut microbiome [13] or of absorption.

**Fig 6.**
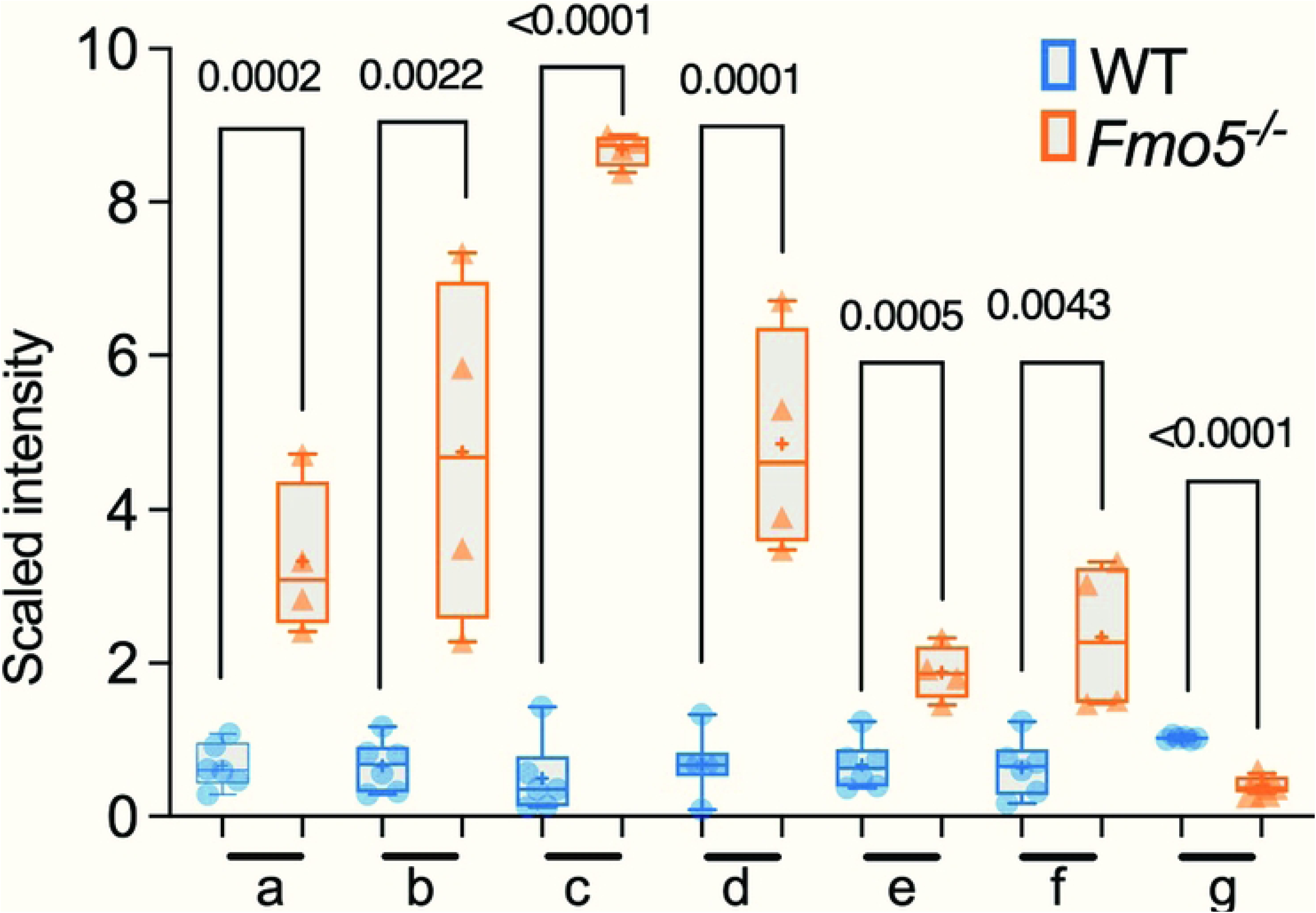
Differences in abundance between *Fmo5^-/-^* and WT mice of products of microbial and/or host-conjugation reactions. a. phenol sulfate; b. phenol glucuronide; c. p-cresol sulfate; d. p-cresol glucuronide; e. 3-indoxyl sulfate; f. trimethylamine *N*-oxide; g. 3-(4-hydroxyphenyl)lactate. + indicates mean value, numbers above box plot are *p* values.

### Upstream regulators

The main upstream regulators predicted by IPA to be affected by disruption of the *Fmo5* gene were the transcription regulators NRF2 and XBP1 and the ligand-dependent nuclear receptors PPARA and PPARG, predicted to be inhibited, and the transcription regulators STAT1 and IRF7, predicted to be activated (Table 7). Consistent with this, expression of genes encoding NRF2 (*Nfe2l2*) was lower and STAT1 and IRF7 was higher in *Fmo5^-/-^* mice (Table 7). There was no significant difference in the expression of *Ppara, Pparg* or *Xbp1*. However, XBP1 is regulated post-transcriptionally. The activities of PPARA and PPARG can be regulated by ligand binding, so FMO5 may act by converting a ligand into an active form or by inactivating an inhibitory ligand.

**Table 7.** Predicted upstream regulators and hepatic expression of their genes in *Fmo5^-/-^* mice compared with WT mice.

| Predicted effect in *Fmo5*<sup>-/-</sup> mice | Protein symbol | Protein name | Z score | *p* value | Percentage change in gene expression (*Fmo5*<sup>-/-</sup> vs WT mice) | padj |
| --- | --- | --- | --- | --- | --- | --- |
| **Inhibited** | NRF2 | Nuclear factor, erythroid 2 like 2 (NRF2) | -5.15 | 2.35e-22 | -19 | 0.033 |
|  | PPARA | Peroxisome proliferator activated receptor alpha | -5.22 | 3.65e-30 | ns | ns |
|  | PPARG | Peroxisome proliferator activated receptor gamma | -4.60 | 2.29e-10 | ns | ns |
|  | XBP1 | X-box binding protein 1 | -6.44 | 3.69e-24 | ns | ns |
| **Activated** | IRF7 | Interferon regulatory factor 7 | +4.12 | 3.33e-7 | +67 | 1.43e-8 |
|  | STAT1 | Signal transducer and activator of transcription 1 | +4.80 | 1.70e-10 | +89 | 0.002 |

ns = not significant

Many of the genes that are downregulated in the liver of *Fmo5^-/-^* mice are activated by NRF2, PPARA, PPARG and/or XBP1, whereas *Stat1*, which is upregulated in *Fmo5^-/-^* mice, is repressed by PPARA (Table 8), and *Parp14, Stat1* and *Irf7*, which are upregulated in *Fmo5^-/-^* mice, are activated by either IRF7 or STAT1 (Table 9).

**Table 8.**
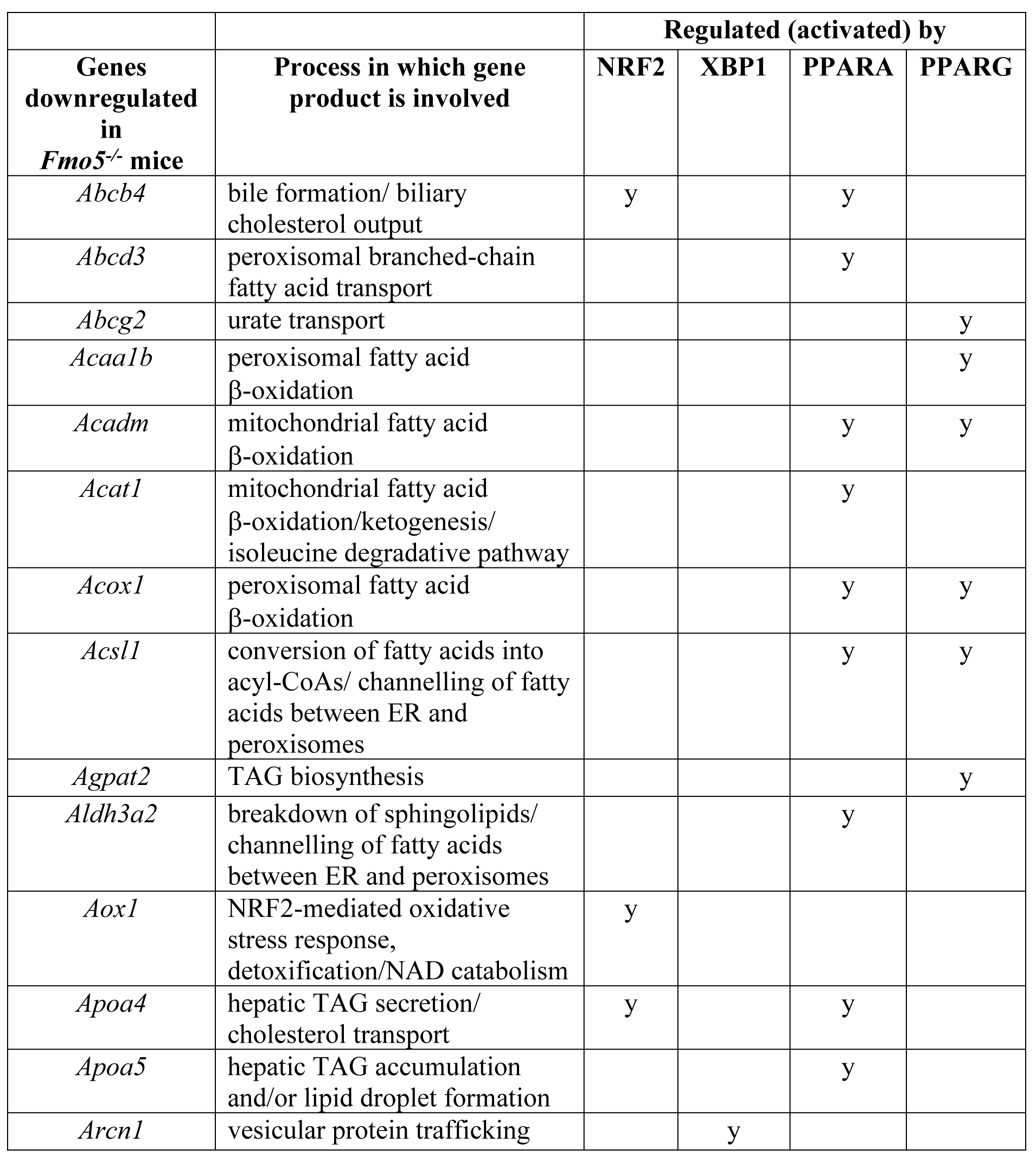
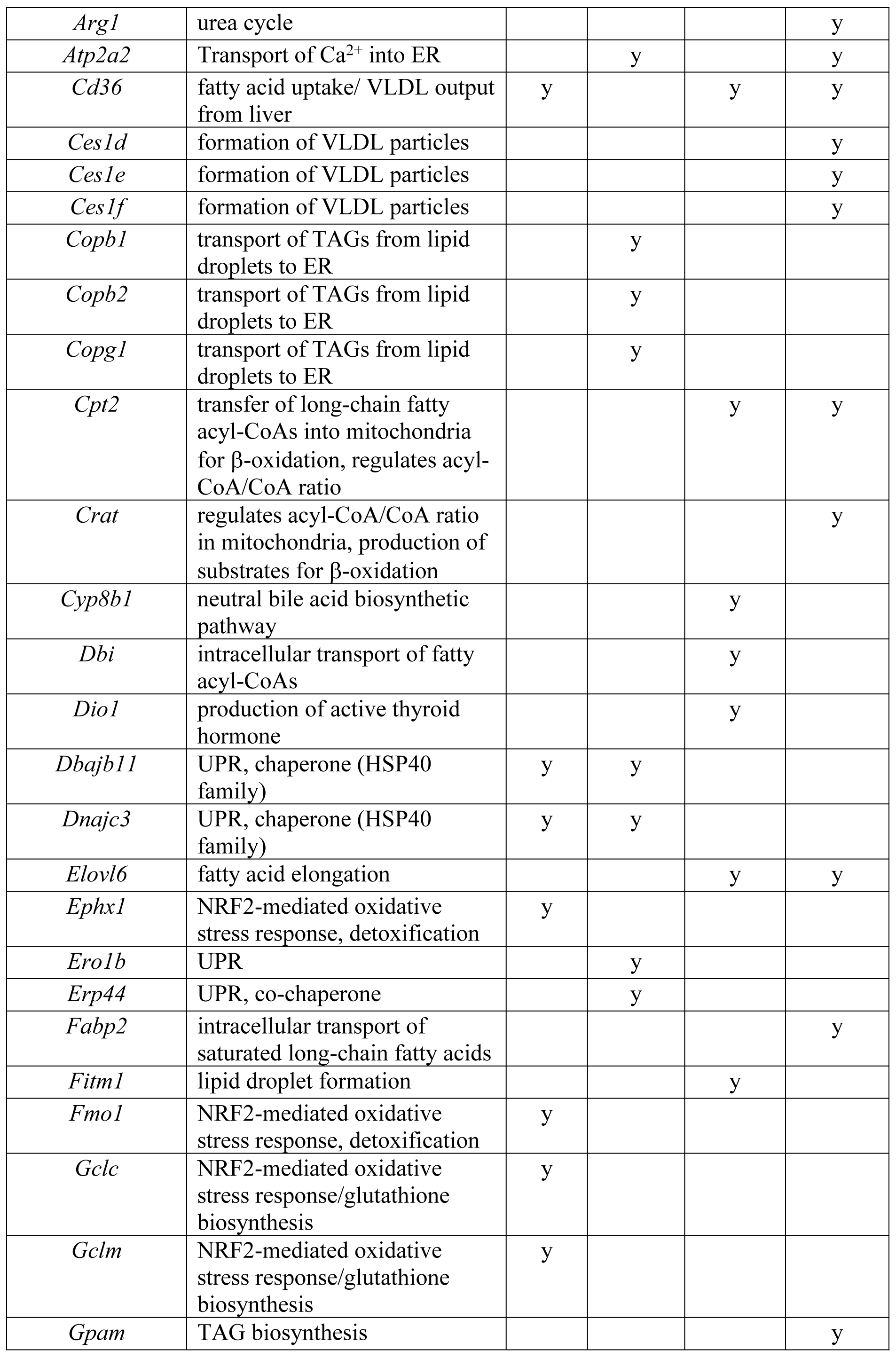
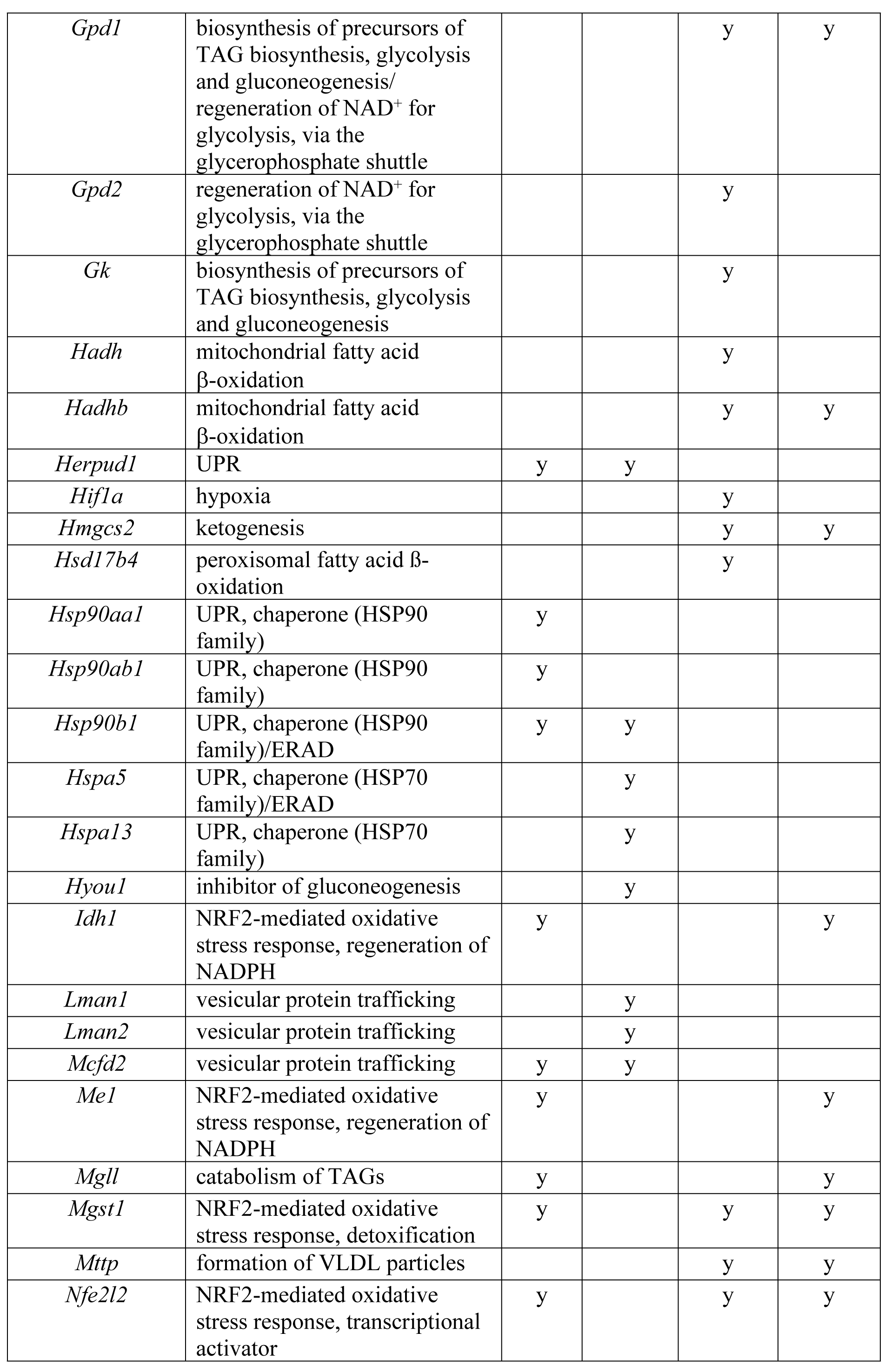
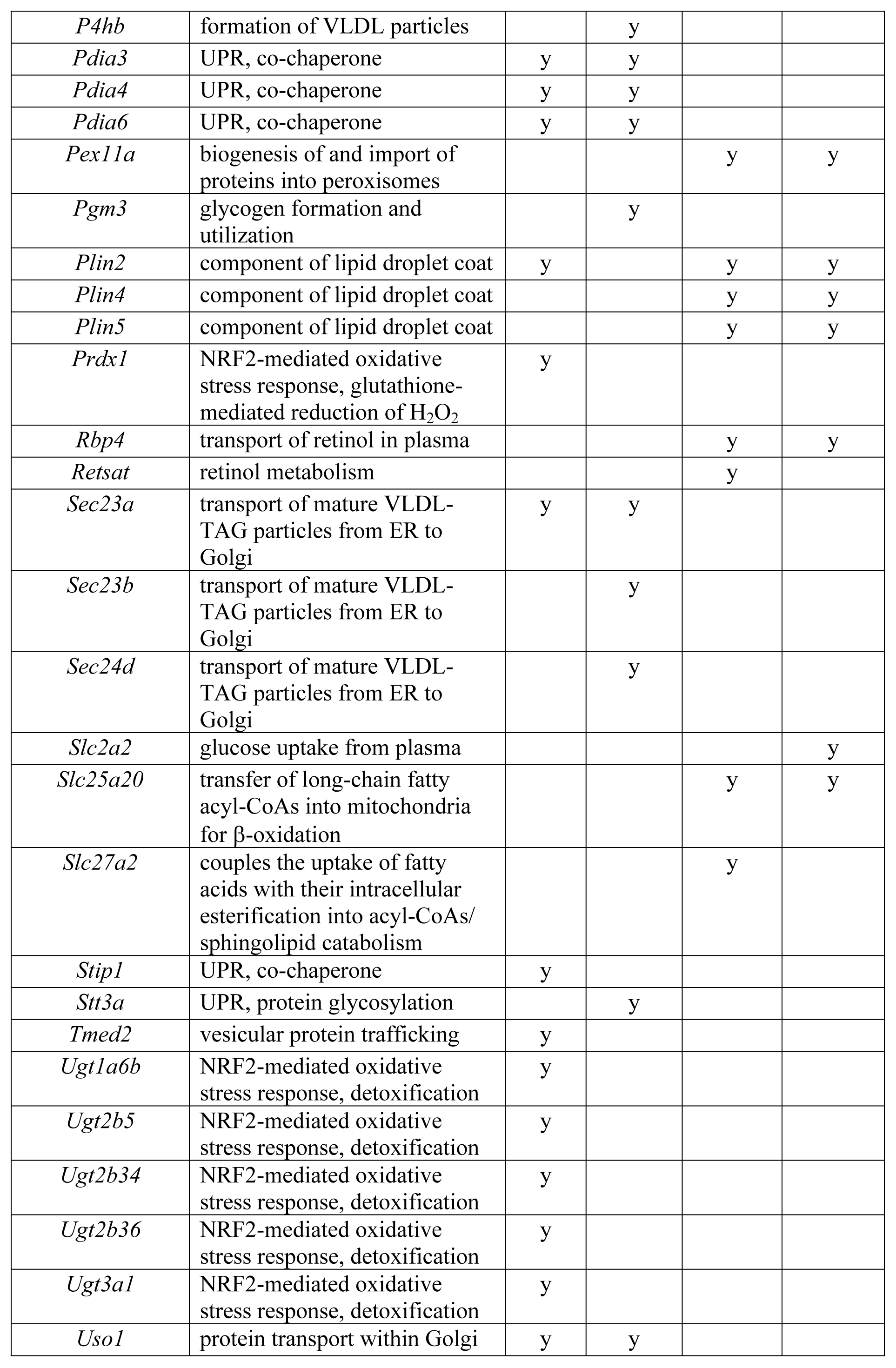
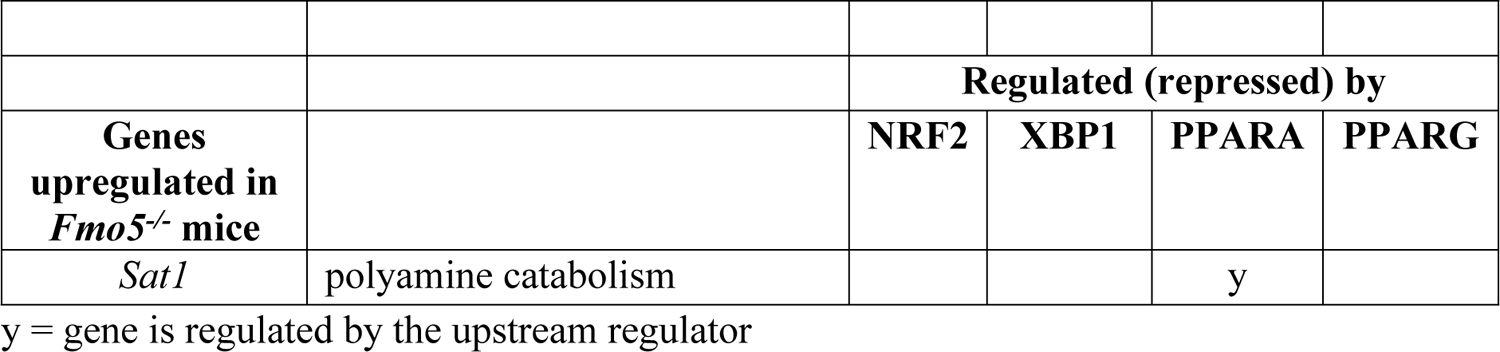
Genes regulated by NRF2, XBP1, PPARA or PPARG that are down- or up-regulated in *Fmo5^-/-^* mice. y = gene is regulated by the upstream regulator.

**Table 9.** Genes regulated by IRF7 and STAT1 that are upregulated in *Fmo5^-/-^*mice.

|  |  |  |  |
| --- | --- | --- | --- |
|  |  | **Regulated (activated) by** |  |
| **Genes upregulated in *Fmo5*<sup>-/-</sup> mice** | **Process in which gene product is involved** | **IRF7** | **STAT1** |
| *Parp14* | mono-ADP ribosylation (MARylation) of proteins | y |  |
| *Stat1* | transcriptional activator | y |  |
| *Irf7* | transcriptional activator |  | y |

y = gene is regulated by the upstream regulator

## Discussion

Metabolomic and transcriptomic approaches reveal that a lack of FMO5 protein has wide-ranging effects on several biochemical pathways and metabolic processes in the liver. The results show that there are marked differences between *Fmo5^-/-^* and WT mice in the abundance of lipids and other metabolites and of mRNAs encoding proteins involved in several metabolic processes, particularly those concerned with the metabolism and transport of lipids, and identify the NRF2*-*mediated oxidative stress response and the UPR as the major downregulated canonical pathways in *Fmo5^-/-^* mice.

NRF2 is a stress-activated transcription factor that orchestrates adaptive responses to oxidative stress by induction of enzymes involved in detoxification, glutathione homeostasis and NADPH regeneration [55]. The expression of *Nfe2l2*, which encodes NRF2, and of several NRF2-activated genes was lower in *Fmo5^-/-^* mice. In addition to its role in oxidative stress, NRF2 modulates intermediary metabolism, in particular fatty acid synthesis and ß-oxidation [55], and several of the phenotypic aspects of *Fmo5^-/-^* mice, lower weight and less fat mass, smaller adipocytes and resistance to diet-induced obesity, are characteristic of *Nfe2l2^-/-^* mice [56], supporting the view that the phenotype of *Fmo5^-/-^* mice is mediated, in part, by decreased expression of *Nfe2l2*.

MAF BZIP transcription factor represses NRF2-activated transcription [19]. Expression of *Maf*, which encodes the transcription factor, is repressed in response to chronic exposure to oxidative stress, glucose and fatty acids [57]. The elevated expression of *Maf* in *Fmo5^-/-^*mice might thus be a consequence of reduced oxidative stress and the lower concentrations of glucose and fatty acids in these animals.

Accumulation of misfolded proteins elicits ER stress. To relieve ER stress, the cell triggers the UPR. Lower expression in *Fmo5^-/-^* mice of *Hspa5* (BiP) and *Hsp90b1*, both of which are induced in response to the presence of misfolded proteins in the ER [58], suggests that in these animals there is less protein misfolding than in WT mice.

Inhibition of fatty acid oxidation and NADPH production, both of which were lower in *Fmo5^-/-^* mice, protects hepatocytes from ER stress [59]. ER stress can be elicited by saturated fats [60], VLDL production [61], insulin [62] and obesity [63]. Thus, lower weight and plasma concentration of insulin [12,13], together with lower abundance of saturated fats in liver and the apparent diminished production of VLDL, would be expected to contribute to reducing ER stress in *Fmo5^-/-^* mice and, thus, attenuating the UPR.

Many of the genes encoding proteins of the UPR that are downregulated in *Fmo5^-/-^* mice are activated by NRF2. The downregulation of *Nfe2l2*, which encodes NRF2, in *Fmo5^-/-^* mice suggests that attenuation of the UPR in these animals is mediated, at least in part, by the effect that disruption of the *Fmo5* gene has on expression of *Nfe2l2*.

The UPR plays a role in regulating lipid metabolism [64], in particular by activating expression of genes encoding proteins involved in ß-oxidation of fatty acids, TAG biosynthesis and formation of VLDL particles. The expression of *Cpt2*, and *Acox1*, involved in fatty acid ß-oxidation, and *Mttp* and *Pdi3, 4* and *6*, involved in VLDL formation, is activated by the UPR. All of these genes were downregulated in *Fmo5^-/-^*mice. Thus, downregulation of fatty acid ß-oxidation and VLDL formation in *Fmo5^-/-^*mice may, in part, be a consequence of attenuation of the UPR.

The down-regulation in the liver of *Fmo5^-/^*mice of expression of genes encoding proteins involved in the UPR (Table 3), including *Hspa5*, which encodes the key UPR protein BiP, is reminiscent of that observed in the livers of mice treated with the stress-inducing molecule tunicamycin [65]. However, a lack of FMO5 does not appear to cause any significant adverse health effects and *Fmo5^-/-^*mice seem well adapted to survive the changes in expression of UPR genes. Interestingly, in a study of the protection by the flavinoid quercetin against mouse liver injury induced by the pyrrolizidine alkaloid clivorine, FMO5 and HSPA5 (BiP) mRNAs were identified in a mouse stress-test array as among the most highly induced (∼100-fold) by quercetin [66] and quercetin treatment reversed the clivorine-induced decrease in the abundance of FMO5 mRNA.

The single FMO of yeast has been shown to oxidize glutathione to glutathione disulfide (GSSG) and is required for correct folding in the ER of proteins containing disulfide bonds [67]. Yeast FMO is induced by the unfolded protein response [68]. In contrast, deletion of mouse *Fmo5* results in an increase in GSSG and decreased expression of genes coding for many components of protein folding, indicating that FMO5 does not generate oxidizing equivalents, but acts upstream of the UPR to promote its activation. The effect of FMO5 may be indirect, for instance, by production of H_2_O_2_ [69], which could stimulate the UPR. Thus, although FMOs are involved in protein folding in both yeast and mammals, the mechanisms by which the enzymes act in these organisms are different.

In addition to downregulation of expression of genes of the NRF2*-*mediated oxidative stress response and the UPR, *Fmo5^-/-^* mice exhibit downregulation of genes involved in hypoxia and cellular stress, suggesting that *Fmo5^-/-^* mice are subject to less oxidative and metabolic stress. FMO5 is considered to be an anti-oxidant protein. However, if amounts of substrate are limiting the catalytic cycle of FMO5 can uncouple and the enzyme can function as an NADPH oxidase, producing H_2_O_2_ [69], thus contributing to oxidative stress.

Our results indicate that several metabolic processes involved in the metabolism and transport of lipids are downregulated in *Fmo5^-/-^*mice. These include the uptake, intracellular transport, elongation and ß-oxidation of fatty acids; ketogenesis; the synthesis and storage of TAG; and the formation and secretion of VLDL particles and cholesterol. Diminished TAG synthesis and storage and formation and secretion of VLDL particles would have a negative effect on the secretion of both TAGs and cholesterol from liver into plasma and their transport to peripheral tissues, thus contributing to the lower plasma cholesterol concentration, the decreased WAT fat stores and, consequently, reduced weight gain of *Fmo5^-/-^*mice [12,13].

Diminished formation of VLDL-particles would also have a negative effect on the transport of plasmalogens from liver into plasma. Lipid rafts are enriched in plasmalogens and cholesterol. The elevated hepatic concentrations of both substances in *Fmo5^-/-^* mice suggest an increase in the abundance of lipid rafts in the liver of these animals.

Increased hepatic concentration of plasmalogens, which are endogenous antioxidants that are preferentially oxidized in comparison with other glycerophospholipids, might protect membrane lipids from oxidation. Increased plasmalogen concentrations have been shown to reduce the accumulation of ROS and protect against hypoxia [70]. Alternatively, increased plasmalogen concentrations might be a consequence of a decrease in ROS. Plasmalogens also protect against hepatic steatosis and steatohepatitis [71].

The hepatokine ANGPTL3 selectively inhibits lipoprotein lipase activity in oxidative tissues such as muscle, routing VLDL-TAG post-prandially for storage in WAT [72]. In addition, it increases plasma concentrations of HDL-cholesterol, via suppression of endothelial lipase [73], and LDL-cholesterol, via an unknown mechanism, and its deficiency is associated with decreased plasma concentrations of glucose and insulin and decreased insulin resistance [74]. Thus, the lower expression of *Angpt3* in *Fmo5^-/-^* mice is likely to contribute to the lower plasma concentrations of cholesterol, glucose and insulin [12,13], the increased glucose tolerance and insulin sensitivity [13] and the smaller amounts of WAT of these animals [12].

ARTD8 and ARTD10, which are encoded by *Parp14* and *Parp10* respectively, catalyze MARylation of proteins. Silencing of *Parp10* increase fatty acid oxidation and glycolysis, rendering cells hypermetabolic [42]. Thus, the increased expression of *Parp10* in *Fmo5^-/-^* mice might contribute to attenuation of fatty acid oxidation and glycolysis in these animals. ARTD8 and ARTD10 also act as RNA-binding proteins and have been implicated, via their role in DNA damage repair, in the response to genotoxic stress [75]. In addition, ARTD10 interferes with NF-κB signalling [76].

A major use of SAM is in the methylation of guanidinoacetate to produce creatine, which consumes about 40% of SAM [77]. The hepatic concentrations of both guanidinoacetate and creatine were higher in *Fmo5^-/-^* mice. Increased consumption of SAM, for production of creatine, may drain the pool of SAM available for methylation of other substrates, with a potential negative effect on epigenetic methylation. The perturbation of one-carbon metabolism in *Fmo5^-/-^* mice could affect age-related DNA hypermethylation [78] and, thus, provide a basis for the delayed metabolic aging of these animals [12]. This is supported by increased hepatic concentrations in *Fmo5^-/-^* mice of a number of methylated products, including several methylated amino acids (S1 Table). In the nematode *C. elegans* it was shown that an FMO modified one-carbon metabolism, which regulated stress resistance and longevity [79].

The expression of *Sat1*, encoding SSAT, which catalyzes the rate-limiting step in polyamine catabolism, was higher in *Fmo5^-/-^* mice. In mice, overexpression of *Sat1* leads to enhanced flux through the polyamine pathway, resulting in increased basal metabolic rate, decreased WAT, improved glucose tolerance and insulin sensitivity, a lean phenotype and resistance to weight gain on a high-fat diet [80,81], all of which are characteristic of *Fmo5^-/-^* mice [12,13].

In addition to *Nef2l2*, mouse knockouts of other genes that are downregulated in the liver of *Fmo5^-/-^*mice share some of the phenotypic characteristics of *Fmo5^-/-^*mice [12,13]. *Cyp8b1^-/-^* mice are resistant to diet-induced obesity and have increased energy expenditure, glucose tolerance and insulin sensitivity [82]; mice deficient in *Aldh1a1* are resistant to diet-induced obesity, have lower fasting plasma glucose concentration and enhanced oxygen consumption [83,84]; *Thrsp^-/-^* mice have a reduced rate of weight gain, less fat mass, and increased glucose tolerance and insulin sensitivity [85]; and THRSP promotes hepatic lipogenesis [86]. However, mice with a liver-specific knockout of *Ide* exhibit increased insulin resistance and glucose intolerance [87].

Our results indicate that disruption of the *Fmo5* gene decreases expression of genes involved in transport and metabolism of the nuclear receptor ligands retinoic acid and thyroid hormone, including *Rbp4*, *Retsat* and *Dio1*. Increased plasma concentrations of RBP4 are correlated with obesity and insulin resistance [88,89], and hepatic depletion of *Retsat* decreases glycolytic flux and de novo lipogenesis [90], consistent with decreased expression of these genes contributing to the metabolic phenotype of *Fmo5^-/-^*mice.

Thyroid hormone induces transcription via interaction with a thyroid hormone receptor-retinoid X receptor (TR-RXR) dimer. However, in the absence of thyroid hormone the dimer represses transcription. Decreased production of T3, as a consequence of lower expression of *Dio1* in *Fmo5^-/-^*mice, might enhance TR-RXR-mediated repression of transcription and, thus, account for the lower expression in *Fmo5^-/-^*mice of the thyroid-hormone inducible genes *Me1* and *Thrsp*.

### Conclusions

Metabolomic and transcriptomic analyses show that disruption of the *Fmo5* gene has wide-ranging effects on the abundance of metabolites and expression of genes in the liver, providing a metabolic basis for the lean phenotype of *Fmo5^-/-^*mice. The results reveal that FMO5 is involved in upregulating the NRF2-mediated oxidative stress response, the UPR and response to hypoxia and cellular stress, indicating a role for the enzyme in adaptation to oxidative and metabolic stress. FMO5 also impacts a wide range of metabolic pathways and processes, principally ones involved in the regulation of lipid homeostasis, the uptake and metabolism of glucose, the generation of cytosolic NADPH, and in one-carbon metabolism.

Although the precise molecular mechanisms by which FMO5 exerts its effects are unknown, our results predict that FMO5 stimulates NRF2, XBP1, PPARA and PPARG regulatory pathways, while inhibiting STAT1 and IRF7 pathways.

## Materials and Methods

### Animal maintenance and sample collection

Male WT C57BL/6J and *Fmo5^-/-^* mice, generated on a C57BL/6J background [12], were bred at University College London. Mice were fed a standard chow diet (Teklad Global Rodent Diet 2019, Harlan Laboratories, Inc., Madison, WI, USA) and housed in adjacent cages with free access to food and water. Blood and tissue samples were collected between 9:00 am and 12 noon. Liver samples were immediately frozen on solid CO_2_ and stored at −80°C. All animal procedures were approved by the local ethics committee (Animal Welfare and Ethical Review Body) and carried out under Home Office Licences in accordance with the UK Animal Scientific Procedures Act.

### Metabolomics

Liver samples (*Fmo5^-/-^*, n= 4 and WT n=6) mice from 32-week-old mice were shipped on solid CO_2_ to Metabolon (Morrisville, NC, USA) for global liver biochemical profiling. The Metabolon platform utilised Ultrahigh Performance Liquid Chromatography-Tandem Mass Spectroscopy (UPLC-MS/MS). Namely, a Waters ACQUITY ultra-performance liquid chromatography (UPLC) and a Thermo Scientific Q-Exactive high resolution/accurate mass spectrometer interfaced with a heated electrospray ionization (HESI-II) source and Orbitrap mass analyzer operated at 35,000 mass resolution. Following log transformation and imputation of missing values, if any, with the minimum observed value for each compound, Welch’s two-sample t-test was used to identify biochemicals that differed significantly (*p* ≤ 0.05) in abundance between *Fmo5^-/-^* and WT mice. The biochemicals identified are given in S1 Table. A volcano plot and boxplots were generated using Prism software, version 9.5.0 (GraphPad Software, Inc., La Jolla, CA). Individual *p* values are stated on each boxplot.

### Transcriptomics

Liver samples (*Fmo5^-/-^*, n= 3 and WT n=3) were shipped on solid CO_2_ to LGC Genomics Gmbh (Berlin, Germany). RNA isolation, cDNA library construction, quality controls and sequencing (NextSeq 500 (1 x 75bp) 400M single reads) were carried out. Demultiplexing of all libraries for each sequencing lane used Illumina bcl2fastq2.17.1.14 software (RAW). Adapter remnants were clipped from all raw reads. Sequencing reads were mapped to mm10 genome using STAR aligner (version 2.6.0c) [91]. Gene-based counting was performed using the STAR package. Differential expression analysis was performed using DESeq2 Bioconductor package (DESeq2_1.22.2) in R [92]. A list of differentially expressed genes was obtained along with Log2-fold change over the control sample (*Fmo5^-/-^*/WT), with an adjusted *p* value (padj) for each gene. The GEO accession number for the RNA sequencing data is GSE180575. Canonical pathways and functions that differed between *Fmo5^-/-^*and WT mice were identified using QIAGEN Ingenuity Pathway Analysis (IPA) (QIAGEN Inc., https://digitalinsights.qiagen.com/IPA) [93]. All data were also subject to manual inspection of function and biochemical pathways and by using Gene Ontology resources [94,95]. A volcano plot was generated using Prism software, version 9.5.0.

### Plasma analysis

Blood was collected and plasma isolated and analysed as described previously [12,96]. Data were analysed with an unpaired Student’s t test (statistical significance defined as a *p* value of <0.05). Statistical analysis was carried out using Prism software (version 9.5). Plasma metabolite concentrations are given in S2 Table.

## Supporting Information

**S1 Table. Biochemicals detected in *Fmo5-/-* and wild-type mice.** Results are expressed as changes of Fmo5-/- vs wild-type mice. Orange = increase; green = decrease.

**S2 Table. Plasma metabolite concentrations of 32-week-old *Fmo5-/-* and WT mice.**

